# Trem2 deficiency attenuates breast cancer tumor growth in lean, but not obese or weight loss, mice and is associated with alterations of clonal T cell populations

**DOI:** 10.1101/2024.09.25.614811

**Authors:** Elysa W. Pierro, Matthew A. Cottam, Hanbing An, Brian D. Lehmann, Jennifer A. Pietenpol, Kathryn E. Wellen, Liza Makowski, Jeffrey C. Rathmell, Barbara Fingleton, Alyssa H. Hasty

**Author notes:** Co-corresponding authors Alyssa H. Hasty, Ph.D. UT Southwestern Medical Center 5323 Harry Hines Blvd., B5.102 Dallas, TX 75390-9003 Barbara Fingleton, Ph.D. 2220 Pierce Ave Nashville, TN 37232-6840. **Conflict of interest disclosure statement:** JCR is a founder and consultant for Sitryx Therapeutics.

## Abstract

Obesity is an established risk factor for breast cancer development and worsened prognosis; however, the mechanisms for this association − and the potential benefits of weight loss − have not been fully explored. The adipose environment surrounding breast tumors, which is inflamed in obesity, has been implicated in tumor progression. An emerging therapeutic target for cancer is TREM2, a transmembrane receptor of the immunoglobulin superfamily that is expressed on macrophages in adipose tissue and tumors. We utilized genetic loss of function (*Trem2*^+/+^ and *Trem*2^-/-^) models and dietary (lean, obese, and weight loss) intervention approaches to examine impacts on postmenopausal breast cancer. Remarkably, *Trem2* deficiency ameliorated tumor growth in lean, but not obese or weight loss mice. Single-cell RNA sequencing, in conjunction with VDJ sequencing of tumor and tumor-adjacent mammary adipose tissue (mAT^Tum-adj^) immune cells, revealed that tumors of lean *Trem2*^-/-^ mice exhibited a shift in clonal CD8^+^ T cells from an exhausted to an effector memory state, accompanied with increased clonality of CD4^+^ Th1 cells, that was not observed in any other diet-genotype group. Notably, identical T cell clonotypes were identified in the tumor and mAT^Tum-adj^ of the same mouse. Finally, an immune checkpoint study demonstrated that αPD-1 therapy restricted tumor growth in lean and weight loss, but not obese mice. We conclude that weight history is relevant when considering potential efficacy of TREM2 inhibition in postmenopausal breast cancer. This work reveals immunological interactions between tumors and surrounding adipose tissue, highlighting significant differences under obese and weight loss conditions.

## Introduction

Obesity is a steadily growing epidemic worldwide^1^ and is associated with increased risk for 13 cancer types, including postmenopausal breast cancer^2^. In the United States, 30.5% of the population were obese (BMI ≥ 30 kg/m^2^) in 2000, which increased to 42.4% in 2018, while severe obesity (BMI ≥ 40 kg/m^2^) rose from 4.7% to 9.2% over the same time period^1^. In addition to an increased risk of postmenopausal breast cancer, epidemiological data also indicate obesity is associated with advanced stage at the time of diagnosis^3^, higher incidence of metastasis^4^, and worse clinical outcomes for obese breast cancer patients regardless of menopausal status^5–7^. The mechanisms linking obesity with increased risk, worsened prognosis, and increased recurrence of breast cancer are not fully understood; however, some key players that have been investigated include increased estrogen, insulin resistance and hyperinsulinemia, increased secretion of adipokines, and heightened inflammation^8^.

Obesity-associated inflammation includes recruitment of macrophages as well as CD8^+^ T cells into abdominal white adipose tissue depots^9,10^. A novel subset of macrophages termed “lipid associated macrophages” (LAMs), with high expression of triggering receptor expressed on myeloid cells 2 (TREM2), have been described predominantly in the visceral adipose tissue of obese mice^11^ and humans^12^. TREM2-expressing LAMs were identified in omental adipose tissue samples from obese individuals, with their proportion strongly correlating with BMI^12^. TREM2 is a transmembrane receptor of the immunoglobulin superfamily that binds a plethora of ligands including bacterial products, DNA, lipoproteins, and phospholipids^13^. As a result of recent discoveries placing TREM2 in a central role of immune signaling, it has emerged as a therapeutic target in Alzheimer’s disease, obesity-related metabolic disease, and cancer^14^.

TREM2 is commonly expressed on tumor associated macrophages (TAMs). For example, carcinomas from various primary sites including skin, liver, lung, breast, bladder, and colon contain TREM2^+^ macrophages^15^. Preclinical models suggest TREM2 expression contributes to an immunosuppressive tumor microenvironment^16^. Studies encompassing subcutaneous models of sarcoma and colorectal cancer as well as orthotopic breast and ovarian cancer models have shown tumor growth attenuation in TREM2-deficient mice and in tumor-bearing mice treated with anti- TREM2 antibody^15,17,18^. However, no work has been done in the postmenopausal breast cancer setting or in the context of obesity and weight loss.

Interestingly, deficiency of TREM2 or treatment with anti-TREM2 antibody augments the efficacy of αPD-1 immune checkpoint therapy^15,18^. The recent discovery of tumor-infiltrating lymphocytes in triple-negative breast cancer (TNBC) has brought to light the immunogenic potential of TNBC in recent years^19^. Pembrolizumab, an αPD-1 therapeutic, and Atezolizumab, an αPD-L1 therapeutic, have been approved by the FDA for treatment of TNBC within defined parameters^20,21^. Work from Makowski’s group demonstrated that αPD-L1 impairs tumor progression in a pre-menopausal breast cancer model in mice that have lost weight through bariatric surgery, but not in obese mice^22^. However, the response of postmenopausal breast cancer to αPD-1 in the settings of obesity or weight loss remains uncertain.

The Women’s Health Initiative Observational Study reported that postmenopausal women with weight loss had a significantly decreased risk of breast cancer compared to postmenopausal women with stable weight^23^. Interestingly, women with sustained weight loss have lower risk of breast cancer compared to women with patterns of weight loss and weight regain^24^. However, whether therapies that modulate immune-tumor cell interactions have altered efficacy in patients who have lost weight is not known. Given the recent data showing that increased inflammatory tone remains unresolved in adipose tissue of mice and humans that have lost weight^25–29^, as well as the dramatic increase in use of potent weight loss drugs, the examination of obesity and weight loss in cancer outcomes is of timely relevance for discovery research.

By implanting lean, obese, and weight loss *Trem2*^+/+^ and *Trem2*^-/-^ mice with murine mammary cancer cells, we discovered that lean, but not obese or weight loss *Trem2*^-/-^ mice were resistant to tumor growth. Accordingly, analysis by single-cell RNA sequencing (scRNAseq) revealed that tumors from lean *Trem2*^-/-^ mice exhibited an increase of highly clonal CD8^+^ and CD4^+^ Th1 cells. Detailed analysis of clonotypes by VDJ sequencing revealed that identical clonotypes were present in both the tumor and tumor-adjacent mammary adipose tissue (mAT^Tum-adj^) of the same mice. Finally, an immune checkpoint inhibitor study demonstrated that αPD-1 restricted tumor growth in the lean and weight loss animals; however, the obese animals were unresponsive to the therapy. We conclude that efficacy of TREM2 modulation and αPD-1 treatment varies under different weight history conditions. Furthermore, weight history in combination with TREM2 modulation impacts the presence and expansion of clonal CD8^+^ T effector memory (CD8^+^ TEM) and CD4^+^ Th1 cells, including matched clones between the tumor and mATTum-adj.

## Methods

### Wild-type mouse studies

Female C57BL/6J mice (JAX; cat. #000664) were used for all wild-type studies. Mice were ovariectomized by Jackson Laboratories at 6 weeks of age and sent to Vanderbilt University. At 8-9 weeks of age, mice were placed on high fat diet (HFD; Research Diets; cat. #D12492) or low fat diet (LFD; Research Diets; cat. #D12450B) to generate lean, obese, and weight loss mice. Lean mice were maintained on the LFD for the entirety of the study. Obese mice were fed LFD for 12 weeks, then switched to HFD for the remaining 11-12 weeks. Contrastingly, the weight loss group was fed HFD for 12 weeks then switched to LFD for the remaining 11-12 weeks. Mice were maintained on 12-hour light-dark cycles. The mouse housing facilities were maintained at 20-25°C and 30-70% humidity. All animal studies were performed only after approval from Vanderbilt’s Animal Care and Use Committee.

### *Trem2*^+/+^ and *Trem2*^-/-^ studies

Wild-type C57BL/6J (JAX; cat. #000664; RRID: IMSR_JAX:000664) and *Trem2* knockout C57BL/6J (JAX; cat. #027197; RRID: IMSR_JAX:027197) mice were purchased from Jackson Laboratories, and colonies were maintained and bred at Vanderbilt University. Mice were first interbred to generate *Trem2* heterozygous mice. *Trem2*^+/+^ and *Trem2*^-/-^ littermate mice resulting from the heterozygous breedings were used in the chow-fed breast cancer study. Subsequently, the *Trem2*^+/+^ and *Trem2*^-/-^ mice resulting from heterozygous breedings were bred independently to generate the quantity of mice necessary for the *Trem2*^+/+^ and *Trem2*^-/-^ diet breast cancer studies. At 6 weeks of age, mice were ovariectomized by the Vanderbilt Mouse Metabolic Phenotyping Center (VMMPC). At 8 weeks of age, mice were placed on either HFD or LFD to generate lean, obese, and weight loss groups as described above.

### Tumor cell implantation

After 20 weeks on diet, mice underwent orthotopic injection of E0771 cells (3 × 10^5^ cells resuspended in PBS (Gibco; cat. #10010-023)) into the fourth left mammary fat pad. Tumors grew for 3-4 weeks as indicated for each study in the Results and Figure Legends. Once palpable, tumors were measured every 1-2 days using electronic calipers to determine tumor volume (V=(L*W*W)/2). The first diet study conducted in wild-type mice were injected with E0771 cells (ATCC; cat. #CRL-3461). All other mice analyzed in this study were injected with E0771 cells that ectopically expressed GFP and PD-L1 (E0771^PD-L1^). The latter cells were generated through a process in which the coding region of murine cd274 was synthesized, codon optimized (Geneart, ThermoFisher) and subcloned into lentiviral vector (pCDH). E0771 cells were infected with the pCDH-PD-L1 expression vector and grown in puromycin to select for cells that stably expressed PD-L1. Mice were sacrificed at day 28 or earlier if volume reached 2000 mm^3^ or ulceration occurred.

### Body composition and glucose tolerance test

Body composition was determined using nuclear magnetic resonance (Bruker Minispec) in the VMMPC. Mice were fasted for 5 h, fasted blood glucose was recorded, and mice were injected intraperitoneally with a glucose bolus (2.0 g dextrose per kg fat free mass)^11^. Blood glucose was measured via tail bleed at 15, 30, 45, 60, 90, 120, and 150 min post-glucose injection with a hand- held glucometer (Bayer Contour Next EZ Meter). Glucose tolerance was analyzed by two-way ANOVA and integrated area under the curve calculated by the trapezoidal rule.

### Magnetic sorting

Magnetic sorting of the adipose tissue stromal vascular fraction was performed using the Miltenyi OctoMACS Separator with MS columns (Miltenyi Biotec; cat. #130-042-201). Magnetic sorting of tumors was performed using the Miltenyi QuadroMACS Separator with LS columns (Miltenyi Biotec; cat. #130-042-401). CD11b (Miltenyi Biotec; cat. #130-093-634) and CD4/CD8 microbeads (Miltenyi Biotec; cat. #130-116-480) and CD8 microbeads (Miltenyi Biotec; cat. #130-116-478) were used according to the manufacturer’s instructions. In brief, whole adipose tissue stromal vascular fraction or single cell suspension of the tumor was stained with microbeads for 15 min at 4°C. Stained sample was run through the magnetic column with subsequent washes. The positive and negative fractions were collected, and the cell pellets were lysed in buffer RLT (Qiagen; cat. #79216) with 1% β-mercaptoethanol and stored at ^-^80°C until used for RNA isolation.

### IgG and αPD-1 administration

Once the average tumor volume of a cohort reached 50-100 mm^3^, intraperitoneal (i.p.) injection with 200 μg/mouse with αPD-1 mAb (clone 29F.1A12; BioXCell; cat. #BE0273; RRID: AB_2687796) or IgG2A isotype control (clone 2A3; BioXCell; cat. #BE0089; RRID: AB_1107769) was initiated. Mice were injected every 2 d for a total of 8 d (total of 4 injections).

### sTREM2 quantification

The protocol for sTREM2 detection in the plasma of mice was adapted from Zhong et al^30^. Briefly, blood was collected at the time of sacrifice via cardiac puncture and centrifuged to yield plasma. A 96-well ELISA plate was coated overnight with TREM2 capture antibody (R&D Systems; cat. #MAB17291; RRID: AB_2208679). The plate was washed 3 times, blocked with 1% BSA for 4 h, and washed another 3 times. Recombinant mouse TREM2 protein (R&D Systems; cat. #1729- T2) was reconstituted according to the manufacturer specifications and diluted to a range of 0-10 ng/mL for the standard curve. Diluted plasma samples and the standard curve were incubated overnight. Biotinylated TREM2 antibody (R&D systems; cat. #BAF1729; RRID: AB_356109) in conjunction with streptavidin HRP (R&D Systems; cat. # DY998) were used to detect sTREM2 at 450 nm on a plate reader.

### Adipose tissue stromal vascular fraction isolation

Mice were sacrificed with isoflurane overdose followed by cervical dislocation and perfusion with 50 mL PBS via the left ventricle. Perigonadal ovarian adipose tissue (oAT) and subcutaneous mAT^Tum-adj^ and contralateral mammary adipose tissue (mAT^Contra^) were dissected, minced with scissors, and digested in 6 mL of 2 mg/mL type IV collagenase (Worthington Biochemical; cat. #LS004189) in PBS with 1% FBS (Gibco; cat. #16140-071) for 30 (oAT) or 45 min (mAT) at 37°C. Samples were vortexed, passed through a 100 μm filter, and centrifuged. Floating adipocytes and supernatant were removed, red blood cells were lysed using ACK Buffer (KD Medical; cat. #RGF-3015), and samples were passed through a 35 μm filter to obtain the stromal vascular fraction as described previously^31^.

### Single cell dissociation of tumor

Tumors were excised from the animal, minced with scissors, and run for one round on the GentleMACS Dissociator Tumor.01 setting in 5 mL 10% FBS 1% P/S RPMI (Gibco; cat. #22400- 089) (cRPMI) in C tubes (Miltenyi Biotec; cat. #130-093-237). Processed samples were incubated in 5 mL cRPMI with 435 U/mL DNase I (Sigma Aldrich; cat. # D5025) and 218 U/mL Collagenase I (Sigma-Aldrich; cat. #C2674) for 30 minutes at 37°C. Digested tumors were vortexed, passed through a 70 μm filter, lysed using ACK buffer (KD medical; cat. #RGF-3015), and passed through a 35 μm filter.

### Single cell RNA sequencing

The CD45^+^ fraction of the mAT^Tum-adj^ and the tumor were isolated for scRNAseq. For the mAT^Tum-adj^, the stromal vascular fraction of the mAT^Tum-adj^ was isolated as described above and stained with DAPI and CD45 (clone 30-F11; Biolegend; cat. #103137). The live CD45^+^ cells were isolated by gating on the DAPI^-^ CD45^+^ fraction on the 4-laser FACSAria III, Bioprotect IV Baker Hood Enclosure. Cells were counted on the Countess III and diluted to 1200 cells/μl. For the tumor, tumors were dissociated to a single cell suspension as described above and counted with the Countess III. Subsequently, 3×10^7^ cells were stained with CD45 microbeads (Miltenyi Biotec; catalogue #130-110-618) and passed first through a LS column on the quadroMACS followed by a second sort of the resulting positive fraction through an MS column on the octoMACS. The CD45^+^ fraction was stained with DAPI and sorted for live cells on the 4-laser FACSAria III, Bioprotect IV Baker Hood Enclosure. Samples were counted with the Countess III and diluted to 1200 cells/μl. The Chromium Next GEM Single Cell 5’ HT v2 kit from 10X Genomics was implemented for both the mAT^Tum-adj^ and tumor samples and run on the Illumina NovaSeq 6000 at the VUMC VANTAGE core facility.

### Single cell RNAseq data processing

Reads from newly generated single cell RNAseq data were processed using Cellranger v.7.2.0 (10X Genomics). For initial quality control of droplets, dropkick v1.2.7^32^ was first performed. Then, dropkick score, mitochondrial RNA content, total number of UMI counts, and feature counts informed thresholding for each individual sample. All cells with > 10% mitochondrial content were filtered. Doublet detection was performed using scDblFinder^33^ and cells identified as doublets were filtered prior to downstream analysis. Count matrices were then loaded into Scanpy v1.9.1^34^ and independent samples were concatenated into one anndata object for tumor specimens and one for adipose specimens. Counts were normalized and log1p transformed. Feature selection for downstream dimensional reduction was performed using the *scanpy.pp.highly_variable_genes* function with *“n_top_genes=5000, flavor=’seurat_v3’”*, and a *batch_key* corresponding to the sample collection batch. For single cell immune receptor sequencing, contig annotations from CellRanger v.7.2.0 were filtered for productive TCRs and imported into anndata objects as metadata.

### Dimensional reduction and data integration

Linear dimensional reduction was performed using principal component analysis with the *scanpy.tl.pca* function with “*svd_solver=’arpack’, n_comps=50”* based on highly variable genes detected by the *scanpy.pp.highly_variable_genes* function. 50 principal components were used for data integration. Concatenated data was integrated with Harmony^35^. For Harmony, the SCANPY implementation was called with the *scanpy.pp.harmony_integrate* function using the sample collection batch and time (morning or afternoon) as categorical covariates. Linear dimensional reduction was performed using Uniform Manifold Approximation and Projection (UMAP) with the *scanpy.tl.umap* function using *“spread=1, min_dist=0.5”*.

### Unsupervised clustering and cell annotation

For the tumor specimens, unsupervised clustering was performed using the Leiden algorithm implemented in the *scanpy.tl.leiden* function with “*resolution=1*”. The *scanpy.tl.rank_genes_groups* function was used to detect cluster-specific enriched genes. A consensus between cluster-specific gene markers and established celltype-specific markers were used to broadly annotate cells. Then, further subclustering of specific populations with various resolution settings was performed to identify higher resolution subclusters. For tumor myeloid cells, the pan-cancer myeloid atlas^36^ was referenced for subpopulation markers where possible. For tumor T cells, ProjectTILs v.3.0^37^ was used with the mouse TIL atlas to predict and annotate subpopulations. For adipose tissue specimens, reference mapping to our previously published murine adipose tissue atlas^10^ was performed using scvi-tools v.1.1.2^38^.

For downstream analyses requiring Seurat objects, scDIOR^39^ was used for object conversion and analyses were performed in R v4.2.2 using Seurat v5.1^40^.

### Differential expression analysis

Differential expression was performed using the Wilcoxon rank sum test in Seurat v5.1 with the *FindMarkers* function using the fast implementation by Presto^41^. Significance was determined by a log fold-change threshold > 0.25 and an adjusted p value of < 0.05.

### RNA isolation, cDNA synthesis, and RT-PCR

Tissues or cells were homogenized and lysed in buffer RLT (Qiagen; cat. #79216) with 1% β- mercaptoethanol. RNA was isolated using the RNeasy Micro Kit (Qiagen; cat. #74004) according to the manufacturer’s instructions. cDNA was synthesized using iScript Reverse Transcription Supermix (Bio-Rad; cat, #1708840). Gene expression was determined using Taqman gene expression assays with FAM-MGB-conjugated primers. All primers were obtained from ThermoFisher Scientific: *Trem2* (cat. #Mm04209422_m1), *Plin2* (cat. #Mm00475794_m1), *Cd36* (cat. #Mm00432403_m1), *Cd9* (cat. #Mm00514275_g1), *Nos2* (cat. #Mm00440485_m1), *Arg1* (cat. #Mm00475988_m1), Itgam (cat. #Mm00434455_m1), *Pdcd1* (cat. #Mm01285676_m1), *Tox* (cat. # Mm00455231_m1), *Tigit* (cat. #Mm03807522_m1), and *Lag3* (cat. #Mm00493071_m1). Experiments were run on the CFX96 Real-Time PCR Detection System (Bio-Rad) as follows: an initial 95°C for 10 min, followed by 40 rounds of 95°C for 10 sec and 55°C for 45 sec with detection at the end of each round. All gene expression was normalized to *Gapdh* and calculated with the -ΔΔCT method.

### Statistical Analysis

Statistical analyses were performed with Prism 10 (GraphPad Software). Data are expressed as mean ± SEM. Student’s *t* test was used to compare two groups. One-way ANOVA was used to compare more than two groups when only one variable was present. Two-way ANOVA was used to compare more than two group when more 2 variables were present. Specific post-hoc tests utilized are indicated within each figure legend. Pearson correlation was used to assess simple linear regression analysis.

### Data availability

Both the raw and processed scRNAseq and VDJ sequencing data in this study have been submitted to the NCBI Gene Expression Omnibus (GEO) and will be made available prior to publication. The GEO Accession Number is **GSE278003**.

## Results

### Models of lean, obese, and weight loss postmenopausal breast cancer

To develop obese and weight loss models of postmenopausal breast cancer, the C57BL/6J mouse strain was selected due to its propensity towards developing obesity and established use in the field of metabolic research (Fig. 1A). At six weeks of age, mice were ovariectomized and recovered on chow diet for two weeks. Subsequently, lean mice were maintained on LFD for the entirety of the study (23-24 weeks), obese mice were fed LFD for 12 weeks then switched to HFD for the remainder of the study (11-12 weeks), and weight loss mice were fed HFD for 12 weeks followed by LFD for the remaining 11-12 weeks (Fig. 1A). Food intake was recorded weekly for all groups (Supp. Fig. 1A & B). Obesity and weight loss were evident by body and oAT mass (Fig. 1B & C), and success of the ovariectomy surgery was determined by uterine atrophy at the end point of the study (Fig. 1D). Glucose tolerance tests were performed after 11 weeks (before the diet switch for the weight loss mice) and 19 weeks on diet. After 11 weeks on HFD, the weight loss group had established obesity and thus exhibited worsened glucose tolerance compared to the mice on LFD (Supp. Fig. 1C). At the 19-week time point, the obese group exhibited worsened glucose tolerance, and the weight loss mice glucose clearance matched the lean controls (Fig. 1E). After 20 weeks on diet, the mice were injected with E0771 cells in the fourth left mammary fat pad and tumors were monitored for 3 weeks. Obese mice exhibited a significant increase in tumor progression measured as volume over time and endpoint tumor mass, while the weight loss mice showed no significant difference from the lean group (Fig. 1F - H).

**Figure 1.**
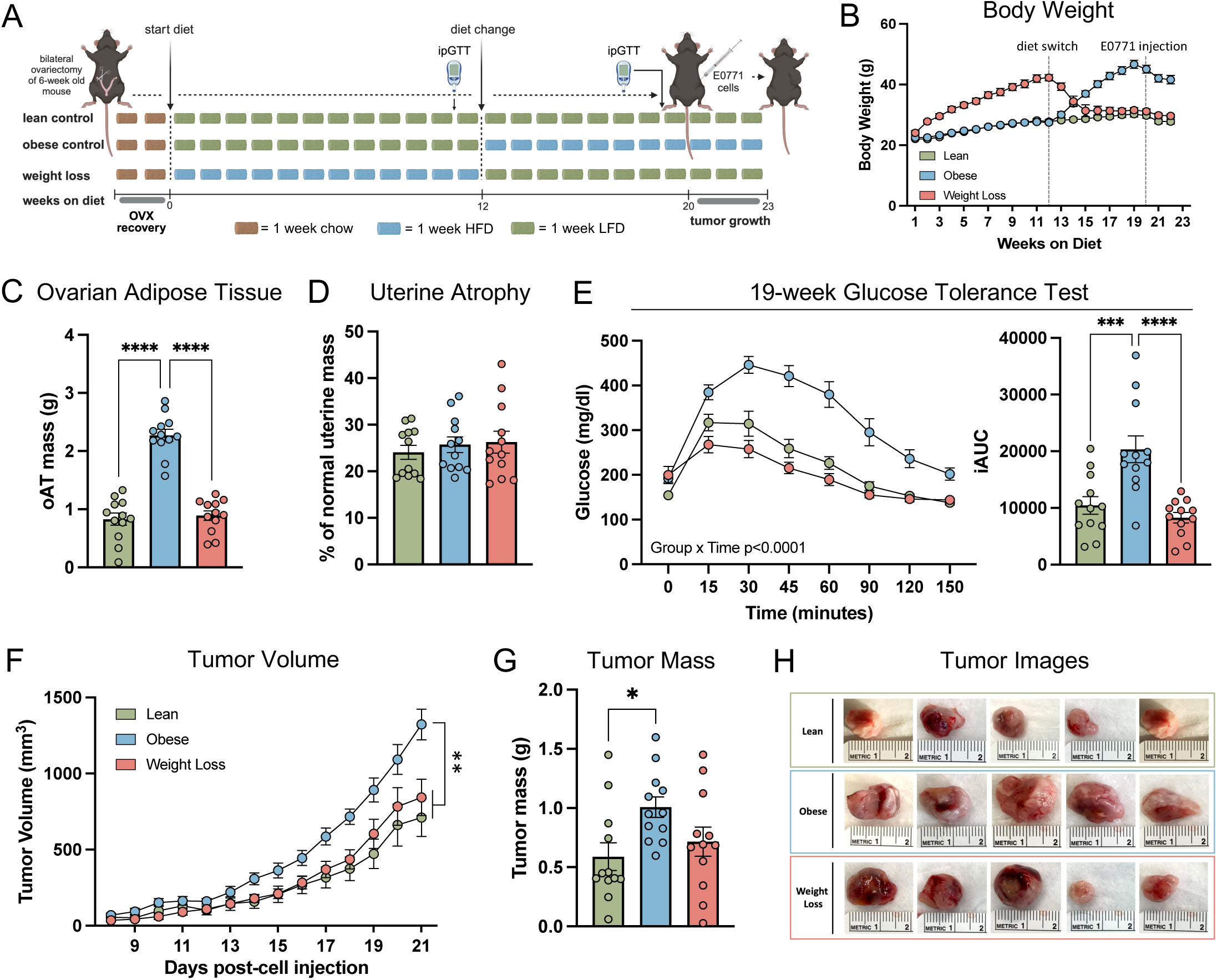
Weight loss rescues tumor growth in postmenopausal breast cancer model. A) Schematic of mouse model of postmenopausal breast cancer in obesity and weight loss including timing of ovariectomy, 10% low fat diet (LFD) and 60% high fat diet (HFD) feeding, intraperitoneal glucose tolerance tests (ipGTT) and orthotopic tumor injections. B) Body weight (g) measured weekly with dashed lines to indicate time of diet switch and E0771 cell injection. For diet groups, green = lean, blue = obese, pink = weight loss. C) Ovarian adipose tissue (oAT) mass at endpoint. D) Uterine atrophy expressed as percent of normal uterine mass. E) Blood glucose (mg/dL) measured over time during 19-week time point ipGTT and corresponding integrated area under the curve (iAUC). F) Tumor volume measured over time. G) End point tumor mass. H) Images of tumors from lean, obese, and weight loss mice. One-way ANOVA with Tukey’s multiple comparisons test was used to compare groups for oAT mass, uterine mass, iAUC from ipGTT, and tumor mass. Repeated measures two-way ANOVA was used for statistical analysis of blood glucose over time in ipGTT and mixed-effects model was used for tumor volume analysis. All data plotted as mean ± SEM for 12 mice per group. *p<0.05, ***p<0.001, ****p<0.0001. Figure 1A was created with Biorender.com.

### Lipid associate macrophage markers were upregulated and positively correlated with tumor mass in tumor-adjacent, but not contralateral, mammary adipose tissue

TREM2 expressing LAMs are associated with obesity^12^ and their immunosuppressive nature has been reported^15,17^. Therefore, expression of LAM genes was quantified in tumors of the lean, obese and weight loss mice. CD11b^+^ cells were magnetically sorted from tumors (Fig. 2A; Supp. Fig. 2A). Expression of LAM genes in the tumor-derived CD11b^+^ cells was not different among dietary groups (Fig. 2B). Next, LAM genes were quantified in mAT^Tum-adj^ and mAT^Contra^ (Fig. 2C). Only *Trem2* expression was increased in the obese mAT^Contra^ compared to the lean mAT^contra^. However, *Trem2* and *Plin2* were increased in obese mAT^Tum-adj^ compared to both the lean mAT^Tum-adj^ and weight loss mAT^Tum-adj^. *Cd36* was increased in obese mAT^Tum-adj^ compared to the lean mAT^Tum-adj^. Furthermore, within the obese and weight loss groups, *Trem2*, *Plin2* and *Cd36* were upregulated in the mAT^Tum-adj^ relative to the mAT^Contra^ (Fig. 2 D - F). To determine whether there is a relationship between tumor growth and LAM gene expression, Pearson correlation analyses between mAT LAM genes and tumor size were performed. There was no correlation between *Trem2*, *Plin2*, or *Cd36* expression and tumor mass in the mAT^Contra^ (Fig. 2G & H; Supp. Fig. 2B); however, *Trem2* and *Plin2* gene expression positively correlated with tumor mass in the mAT^Tum-adj^ (Fig. 2G & H; Supp. Fig. 2C).

**Figure 2.**
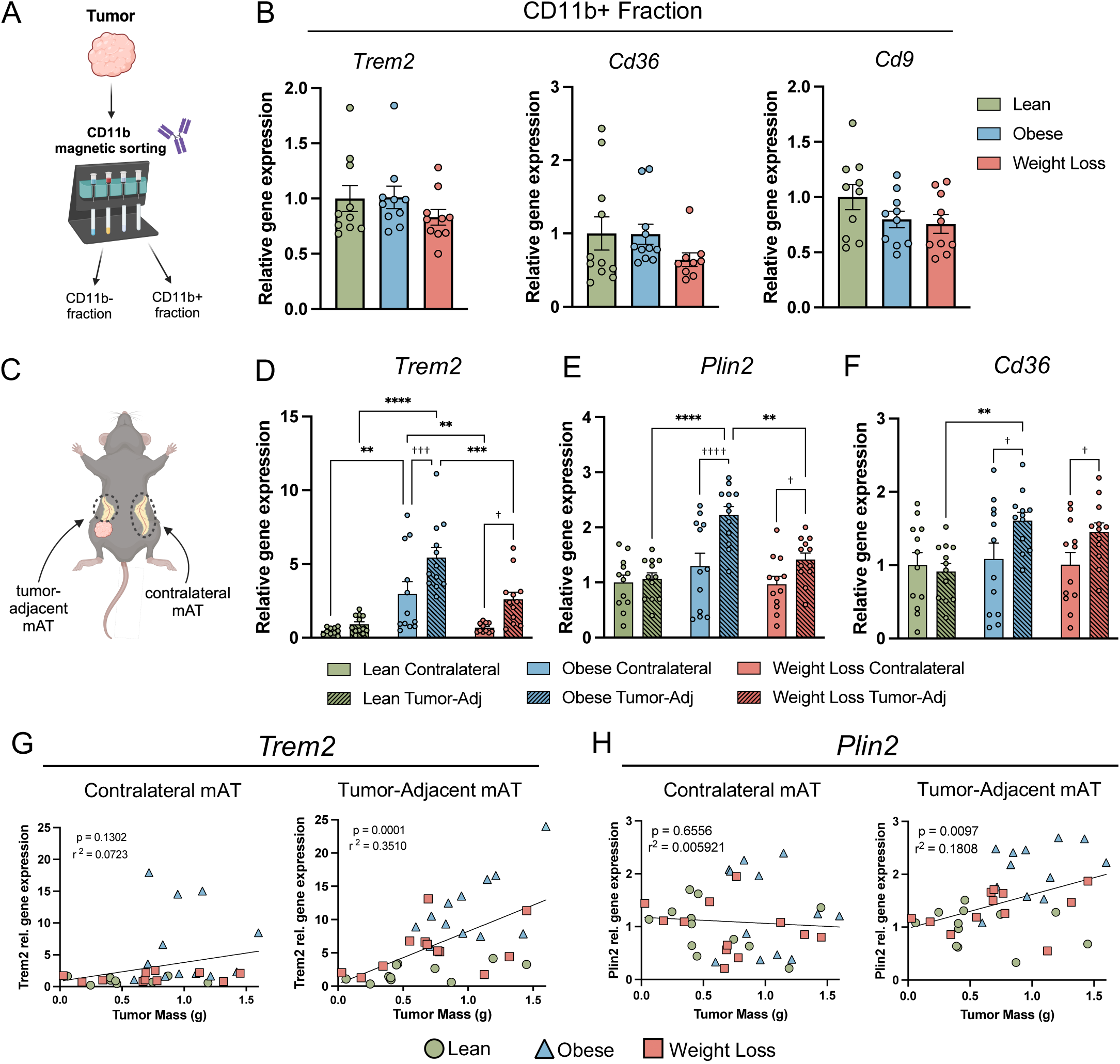
Lipid associated macrophage marker genes are increased in obese tumor-adjacent mammary adipose tissue (mAT^Tum-adj^) and positively correlate with tumor mass. B) Schematic for magnetically sorting CD11b^+^ cells from E0771 tumors using Miltenyi Octomacs. C) *Trem2*, *Cd36*, and *Cd9* relative gene expression by RT-PCR in tumor CD11b^+^ fraction. C) Anatomical schematic of subcutaneous mAT^Tum-adj^ and contralateral mammary adipose tissue (mAT^Contra^). D - F) *Trem2*, *Plin2*, *Cd36* relative gene expression by RT-PCR from whole tissue lysates of mAT^Contra^ and mAT^Tum-adj^. G-H) Correlation analysis of tumor mass vs. *Trem2* and *Plin2* gene expression in mAT^Contra^ and mAT^Tum-adj^ with linear regression fit. Figures D-F, solid bar = mAT^Contra^, hashed bar = mAT^Tum-adj^. One-way ANOVA tests were used to compare tumor LAM marker gene expression among groups. Two-way ANOVA with Tukey’s multiple comparison tests were used to compare groups for all mAT gene expression. Simple linear regression analysis was applied to correlation plots and the p value and coefficient of determination (r^2^) is displayed on each graph. All data plotted as mean ± SEM for 8-12 mice per group. *denotes *diet* effect, †denotes *tissue* effect (mAT^Contra^ vs. mAT^Tum-adj^), **p<0.01, ***p<0.001, ****p<0.0001; †p<0.05, †††p<0.001, ††††p<0.0001. Figures 2A & C were created with Biorender.com.

### Trem2 deficiency decreased tumor growth in postmenopausal breast cancer model

Other groups have demonstrated TREM2 deficiency or treatment with anti-TREM2 monoclonal antibody results in decreased tumor growth in models of sarcoma, colorectal cancer, pre-menopausal breast cancer, and ovarian cancer^15,17,18^. To determine if TREM2 ablation attenuates tumor growth in the postmenopausal breast cancer model, *Trem2*^+/+^ and *Trem2*^-/-^ female C57BL/6J mice were ovariectomized and maintained on a chow diet (Fig. 3A). Plasma soluble TREM2 (sTREM2) was absent in the *Trem2*^-/-^ animals (Fig. 3B). Throughout the duration of the study, no differences in body weight or glucose tolerance (ipGTT at 14-wks) were observed between genotypes (Fig. 3C; Supp. Fig. 3A), and oAT and mAT mass were not different at the study endpoint (Supp. Fig. 3B & C). Twenty-five weeks after ovariectomy, mice were injected with E0771^PD-L1^ cells, and tumors were monitored for 4 weeks. Both tumor progression and tumor mass at endpoint were significantly reduced in the *Trem2*^-/-^ mice compared to *Trem2*^+/+^ controls (Fig. 3D & E; individual tumor growth curves - Supp. Fig. 3D & E). *Cd36* gene expression was significantly reduced in oAT of *Trem2*^-/-^ mice, with a slight reduction of *Plin2* and *Cd9* (Fig. 3F). Both M1-like and M2-like macrophage markers, *Nos2* and *Arg1,* respectively, were decreased in the oAT (Fig. 3F). Previous reports in male epididymal adipose tissue indicated profound adipocyte hypertrophy in Trem2 deficient mice^11,12^. In this model, the *Trem2*^-/-^ mAT^Tum-adj^ exhibited a shift in size from small (25-49μm diameter) adipocytes toward large (70-99μm diameter) adipocytes (Fig. 3G - I), which is the first time this has been reported in female mAT.

**Figure 3.**
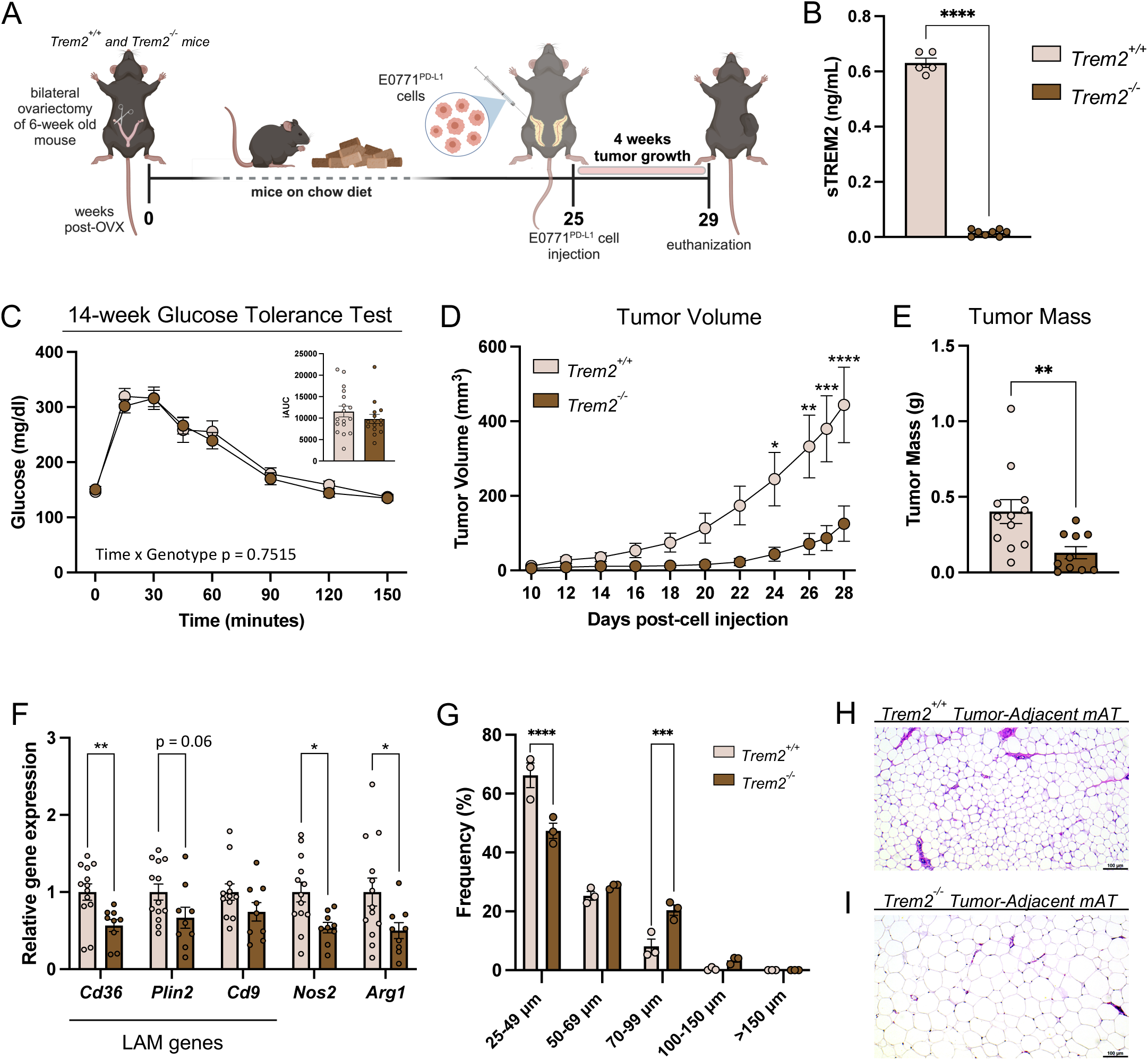
Trem2 ablation attenuates tumor growth, decreases lipid associated macrophages, and increases adipocyte hypertrophy in a chow-fed postmenopausal breast cancer model. A) Schematic of study design for chow-fed postmenopausal breast cancer model in Trem2^+/+^ and Trem2^-/-^ mice. B) sTREM2 (ng/mL) in plasma from female Trem2^+/+^ and Trem2^-/-^ mice measured by ELISA. C) Blood glucose (mg/dL) measured during 14-week time point intra-peritoneal glucose tolerance test (ipGTT) and corresponding integrated area under the curve (iAUC). D) Tumor volume (mm^3^) over time measured every 1-2 days. E) Tumor mass (g) at study endpoint. F) Relative gene expression by RT-PCR of LAM and macrophage markers from oAT whole tissue lysates. G) Adipocyte sizing frequency in tumor-adjacent mammary adipose tissue (mAT^Tum-adj^). Diameter: small = 25-49μm, medium = 50-69μm, large = 70-99μm, very large = 100-150μm, extremely large = >150μm. H-I) Representative image of Trem2^+/+^ and Trem2^-/-^ mAT^Tum-adj^ stained with hematoxylin and eosin. Scale bar represents 100μm. Two-way ANOVA with uncorrected Fisher’s LSD test was used to compare blood glucose over time in the ipGTT. Two-way ANOVA with Sidak’s multiple comparison test was used to compare tumor volumes over time between genotypes. Unpaired two-tailed t-tests were used to compare groups for sTREM2 ELISA, iAUC, tumor mass, LAM gene expression, and each binning diameter size in adipocyte sizing analysis. All data plotted as mean ± SEM for 10-13 mice per group. *denotes *genotype* effect (Trem2^+/+^ vs. Trem2^-/-^) *p<0.05, **p<0.01, ***p<0.001, ****p<0.0001. Figure 4A was created with Biorender.com.

### Trem2 ablation attenuated tumor growth in lean, but not obese or weight loss animals

To assess the impact of TREM2 ablation on tumor growth in the setting of obesity and weight loss, female ovariectomized *Trem2*^+/+^ and *Trem2*^-/-^ mice were placed on LFD and HFD to generate lean, obese, and weight loss groups and injected with E0771^PD-L1^ after 20 weeks on diet. There were no differences in body weight trajectories between genotypes within each diet group (Fig. 4A). Plasma levels of sTREM2 were increased in the obese *Trem2*^+/+^ animals relative to both the lean and weight loss *Trem2*^+/+^ mice, and the *Trem2*^-/-^ animals demonstrated absence of sTREM2 (Fig. 4B). Similar to chow-fed lean animals (Fig 3D & E), tumor growth in the LFD-fed lean *Trem2*^-/-^ animals was attenuated relative to the lean *Trem2*^+/+^ controls as demonstrated by tumor mass at endpoint and tumor volume over time (Fig. 4C & D). However, *Trem2* deficiency did not reduce tumor growth in obese or weight loss mice (Fig. 4C, E & F).

**Figure 4.**
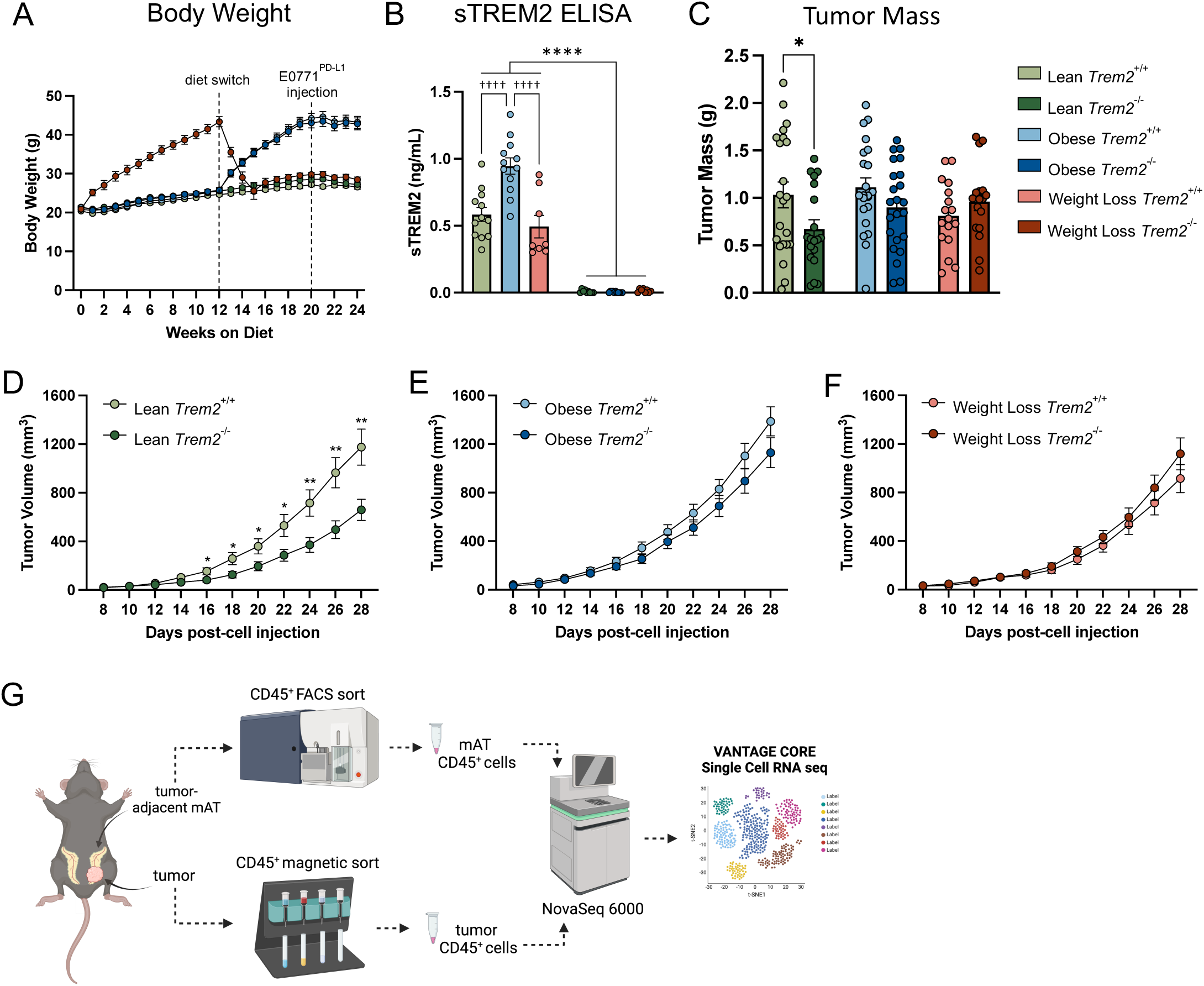
Lean, but not obese or weight loss, Trem2^-/-^ mice exhibit decreased tumor growth. A) Body weight (g) measured weekly with dashed lines to indicate time of diet switch and E0771^PD-L1^ cell injection. For diet groups and genotypes, light green = lean Trem2^+/+^, dark green = lean Trem2^-/-^, light blue = obese Trem2^+/+^, dark blue = obese Trem2^-/-^, light pink = weight loss Trem2^+/+^, and maroon = weight loss Trem2^-/-^. B) sTREM2 (ng/mL) in plasma from Trem2^+/+^ and Trem2^-/-^ mice measured by ELISA.C) Tumor mass (g) at study endpoint. D-F) Tumor volume (mm^3^) over time for lean, obese, and weight loss Trem2^+/+^ and Trem2^-/-^ mice, respectively. G) Diagram of tumor and tumor-adjacent mammary adipose tissue CD45^+^ cell sorting and processing for single cell RNA Sequencing. Mixed effects analysis with uncorrected Fisher’s LSD test was used compare tumor volumes over time between genotypes in the lean group (one missing value in lean Trem2^+/+^ group at day 28). Two-way ANOVA with Sidak’s multiple comparison test was used to compare tumor volumes over time between genotypes in obese and weight loss groups. Two-way ANOVA with uncorrected Fisher’s LSD test was used to compare tumor mass between genotypes in lean, obese, and weight loss groups. *denotes *genotype* effect (Trem2^+/+^ vs. Trem2^-/-^), †denotes *diet* effect, *p<0.05, **p<0.01, ****p<0.0001 for genotype differences (Trem2^+/+^ vs. Trem2^-/-^) and ††††p<0.0001 for diet differences. All data plotted as mean ± SEM for n=4-23 per group. Figure G was created with Biorender.com.

### Single cell RNA sequencing revealed increased expression of antigen presentation genes in tumor-associated macrophages in lean Trem2^-/-^ mice

To gain deeper insights into the immunological landscape of the tumor and the mAT^Tum-adj^, we conducted single-cell RNA sequencing on CD45^+^ immune cells (Fig. 4H). UMAP analysis identified 21 clusters in the tumor and 33 clusters in the mAT^Tum-adj^ (Supp. Fig. 4A & B). The top 3 differentially expressed genes (DEGs) for each cluster are shown in Supplemental Figures 4C and D. Because *Trem2* expression is restricted to myeloid cells, myeloid populations from the tumor and mAT^Tum-adj^ were subset and reclustered (Fig. 5A & B), and the frequencies of each cell population within each tissue were calculated (Fig. 5C & D). Our RT-PCR data showed that whole tumor *Trem2* expression is not impacted by obesity or weight loss (Fig 2B). In our scRNAseq data set, *Trem2*, which is predominantly expressed in the Macro_C1qc subcluster, a more pro-inflammatory macrophage population involved in antigen presentation, shows no variation between diet groups in the tumor (Fig. 5E). However, in mAT^Tum-adj^, obesity increased *Trem2* expression in the LAM population, as has been shown to occur in other adipose tissue depots^10,12^ (Fig. 5F). The top 5 DEGs between the lean *Trem2*^+/+^ and lean *Trem2*^-/-^ were determined within the tumor Macro_C1qc population and subsequently assessed in all groups (Fig. 5G). Four out of the top 5 DEGs, *H2-Eb1*, *H2-Ab1*, *H2-Aa*, and *Cd74*, were relatively increased in the lean *Trem2*^-/-^ Macro_C1qc (tumor) and LAM (mAT) populations compared to the lean *Trem2*^+/+^ group (Fig. 5H). These genes are integral to antigen presentation, which directed our focus to T cell populations.

**Figure 5.**
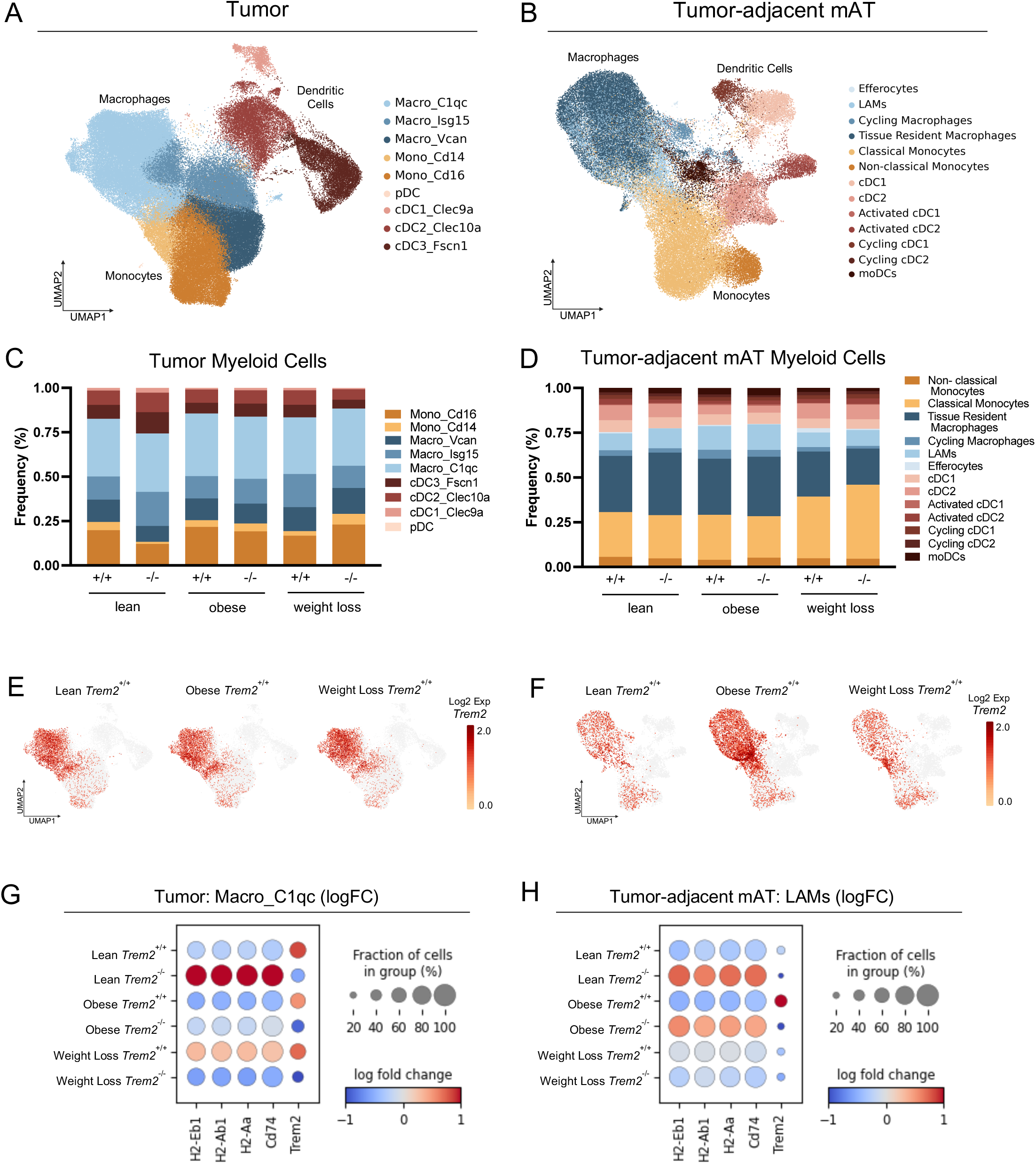
Myeloid populations in tumors and tumor-adjacent mammary adipose tissues (mAT^Tum-adj^) analyzed by single-cell RNA sequencing. A-B) Uniform Manifold Approximation and Projection (UMAP) of tumor (A) and mAT^Tum-adj^ (B) myeloid cell subclusters from merged conditions labeled broadly by cell type category and colored by high-resolution cell type identities. C-D) Myeloid cell subcluster proportions expressed as frequency (%) split by diet-genotype groups for tumor (C) and mAT^Tum-adj^ (D). E-F) *Trem2* expression in lean, obese, and weight loss Trem2^+/+^ mice in tumor (E) and mAT^Tum-adj^ (F). G) Dot plot of the top 5 differentially expressed genes (DEGs) between lean Trem2^+/+^ and lean Trem2^-/-^ tumor Macro_C1qc subclusters. Genes include *H2-Eb1*, *H2-Ab1*, *H2-Aa*, *Cd74*, and *Trem2*. Log fold change is shown for all 6 diet-genotype groups. H) Dot plot of *H2-Eb1*, *H2-Ab1*, *H2-Aa*, *Cd74*, and *Trem2* log fold change in the mAT^Tum-adj^ LAM subcluster shown in all 6 diet-genotype groups.

### Clonal T cell populations were shifted from an exhausted to effector state in tumors from lean but not obese or weight loss Trem2^-/-^ mice

Initial analysis of the tumor T cell populations included both cell proportions, shown as cells/g tumor, and frequency (Fig. 6A & B), as well as subclustering of the tumor T cell populations (Fig. 6C). In the lean groups, *Trem2*^-/-^ tumors showed an increase of CD8^+^ TEM and CD4^+^ Th1 cells calculated as percentage and number of cells/g tumor compared to *Trem2*^+/+^ controls. These two T cell populations also increased in *Trem2*^-/-^ conditions compared to *Trem2*^+/+^ conditions in the obese mice; however, there were markedly reduced cells/g tumor in obese animals of both genotypes. In contrast to the lean mice, in the weight loss group, CD8^+^ TEM and CD4^+^ Th1 cells were reduced with *Trem2* knockout compared to their diet-matched controls (Fig. 6A & B). Because T cell clonal expansion is integral for targeting and killing tumor cells, tumor T cell clonality (number of cells sharing a clonotype) was visualized by projection of clonal cell counts onto the re-clustered tumor T cell UMAP split by diet and genotype (Fig. 6D - I). Highly clonal populations of exhausted CD8^+^ T cells (TEX) were found in lean *Trem2*^+/+^ tumors (Fig. 6D). In tumors from lean *Trem2*^-/-^ mice, the clonal populations were detected in CD8^+^ TEM and CD4^+^ Th1 cells (Fig. 6E). In the obese animals, there were fewer highly clonal populations overall; however, the shift toward increased CD8^+^ TEM populations in the *Trem2*^-/-^ persisted compared to *Trem2*^+/+^ tumors (Fig. 6F & G). In the weight loss animals, the *Trem2*^+/+^ group harbored a robust clonal population of CD8^+^ TEX cells while the weight loss *Trem2*^-/-^ T cells demonstrated a marked decrease of overall clonality (Fig. 6H & I). Strikingly, the rich abundance of highly clonal CD4^+^ Th1 cells in lean *Trem2*^-/-^ tumors was absent from the other diet and genotype groups. Binning of T cells clones into singlets and low, moderate, or high clonality bins (low = x ≤ 12 (median); moderate = 13 ≤ x ≤ 79 (3^rd^ quartile cutoff); high = x >79) reinforced this increase of CD4^+^ Th1 cells in the lean *Trem2*^-/-^ tumors (Fig. 6J). In the lean and obese groups, binning also highlighted the expansion of highly clonal CD8^+^ TEM in the Trem2^-/-^ mice compared to their diet-matched *Trem2*^+/+^ controls; however, this robust expansion was not observed in the weight loss animals. Given the substantial differences in tumor T cell clonality between diet and genotype groups, and considering the impact of the adipose tissue microenvironment, we proceeded to investigate the clonotypes not only within the tumor but also in the mAT^Tum-adj^.

**Figure 6.**
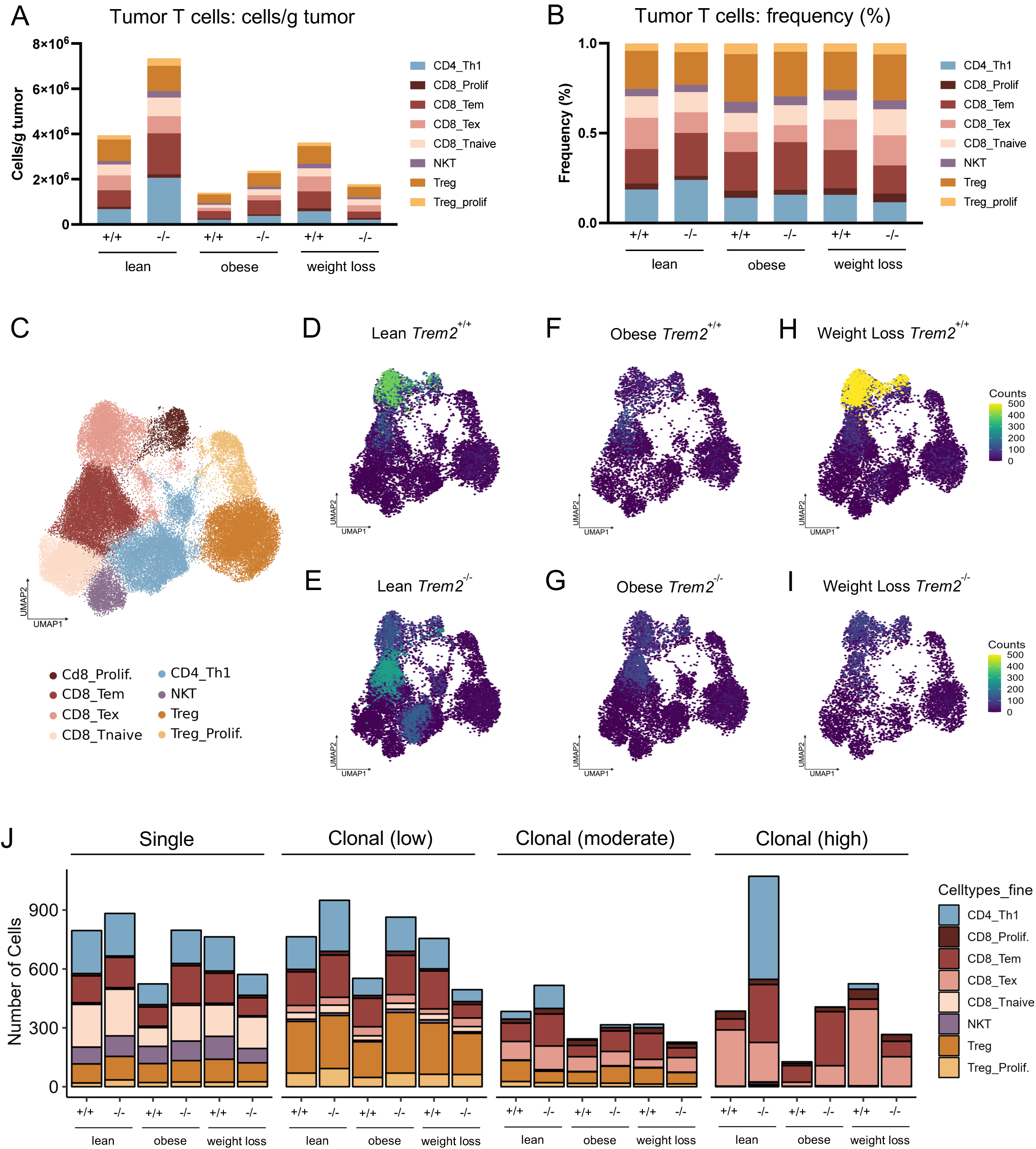
Visualization and binning of Tumor T cell clones reveals expansion of highly clonal CD4^+^ Th1 cells in lean Trem2^-/-^ tumors. A) Stacked bar graph of tumor T cell subclusters indicating cells per gram of tumor across all 6 diet-genotype groups. B) Tumor T cell subcluster proportions expressed as frequency (%) across all 6 diet-genotype groups. C) Uniform Manifold Approximation and Projection (UMAP) of tumor T cell subclusters from merged conditions colored by high-resolution cell type identities. D-I) Tumor T cell clones with a clonal population of two or more visualized by mapping clonal cell counts, indicated by color, onto the tumor T cell UMAP. The data is split as labeled by diet and genotype. J) Stacked bar graph showing the distribution of tumor T cell subclusters, binned as single cells or clonal low, clonal moderate, or clonal high (low = x ≤ 12 (median); moderate = 13 ≤ x ≤ 79 (3^rd^ quartile cutoff); high = x >79) shown for all 6 diet-genotype groups.

### Identification of identical clonotypes in tumor and mAT^Tum-adj^

Clonally expanded T cells have been shown to reside in both tumors and adipose tissue, but whether there are any shared clones that traffic between the two has not been reported. By sequencing the beta chain of the T cell receptor (TCRβ), we determined clonotypes based on the complementarity determining region 3 (CDR3). T cell clones were categorized into three groups: tumor-only clones, which were identified exclusively in the tumor, adipose-only clones, which were found solely in the mAT^Tum-adj^, and dual-tissue clones, which were identified by the presence of the same TCRβ in both the tumor and the mAT^Tum-adj^ (Fig. 7A). Heat maps of tumor-only and adipose-only clones, when overlaid on their respective re-clustered T cell UMAPs, show that single tissue clones were distributed across all identified T cell populations within each diet-genotype group (Fig. 7 B - D). Heat maps of tumor dual-tissue clones revealed that lean *Trem2*^+/+^ control tumors primarily contain CD8^+^ TEM dual-tissue clones, whereas lean *Trem2*^-/-^ tumors exhibit dual-tissue clones distributed across CD8^+^ TEX, CD8^+^ TEM, and CD4^+^ Th1 populations (Fig. 7E). Both obese genotype groups harbor primarily CD8^+^ TEM dual-tissue clones in the tumor (Fig. 7E). However, the weight loss animals exhibit a flipped phenotype from the lean animals wherein the *Trem2*^+/+^ tumors show a spread of dual-tissue clones across CD8^+^ TEX, CD8^+^ TEM, and CD4^+^ Th1 populations and the *Trem2*^-/-^ tumors primarily contain CD8^+^ TEM clones (Fig. 7E). Heat maps of mAT^Tum-adj^ dual-tissue clones show that lean and obese *Trem2*^-/-^ mAT^Tum-adj^ exhibit more dual tissue clones compared to their *Trem2*^+/+^ diet-matched controls; however, both weight loss groups show a diminishment of dual-tissue clones within the mAT^Tum-adj^ (Fig. 7F).

**Figure 7.**
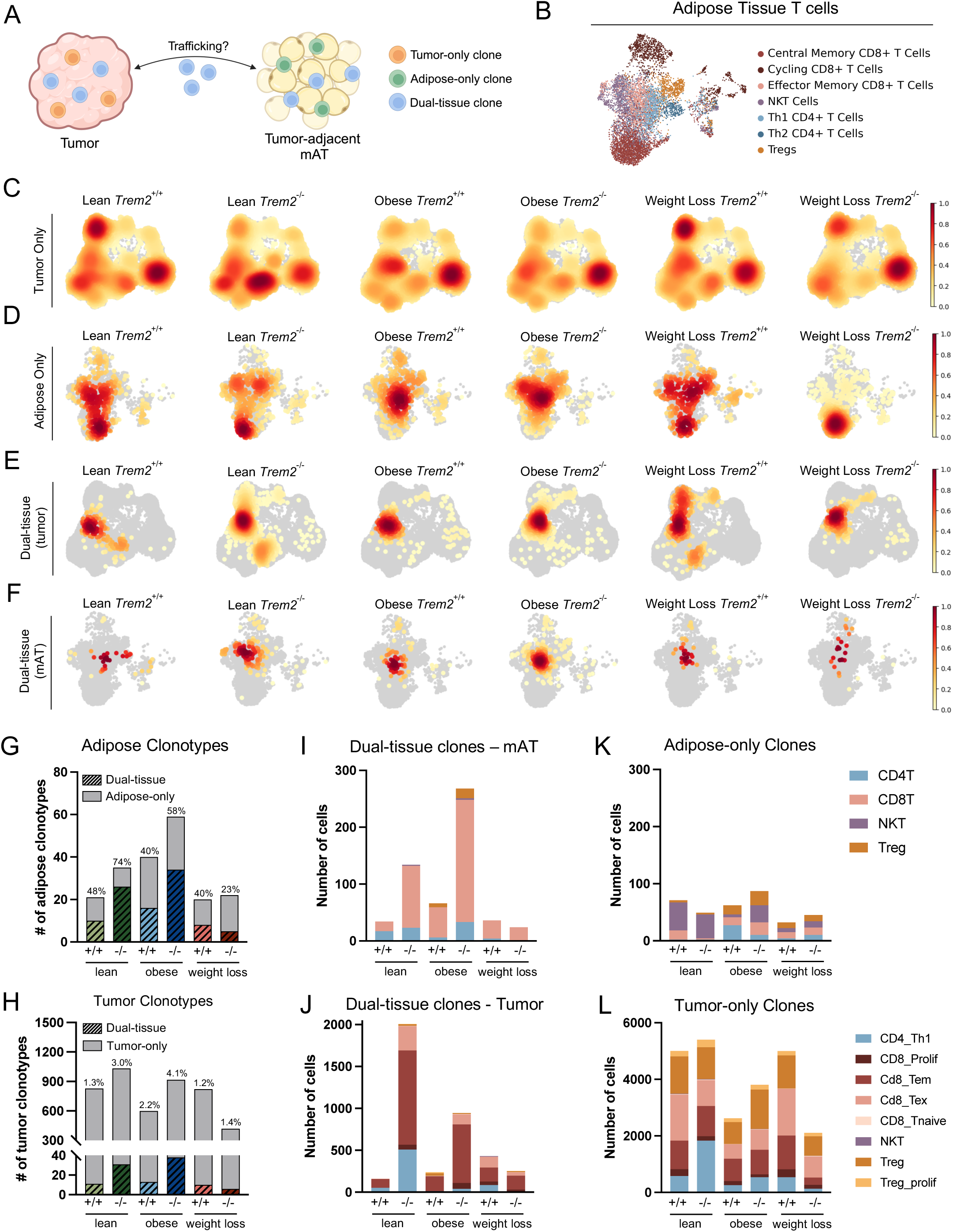
Identification of dual-tissue clones in tumor and tumor-adjacent mammary adipose tissue (mAT^Tum-adj^). A) Schematic illustrating tumor-only, adipose-only, and dual-tissue clones. B) Uniform Manifold Approximation and Projection (UMAP) of mAT^Tum-adj^ T cell subclusters from merged conditions colored by high-resolution cell type identities. C) Heat map of tumor-only clones projected onto tumor T cell UMAP. D) Heat map of adipose-only clones projected on mAT^Tum-adj^ T cell UMAP. E) Heat map of dual-tissue clones identified in the tumor projected on tumor T cell UMAP. F) Heat map of dual-tissue clones identified in the mAT^Tum-adj^ projected on mAT^Tum-adj^ T cell UMAP. G) Stacked bar graph showing the number of clonotypes identified in the mAT^Tum-adj^, sorted by dual-tissue and adipose-only clones. The percentage of dual-tissue clones for each diet-genotype group is displayed at the top of each bar. H) Stacked bar graph showing the number of clonotypes identified in the tumor, sorted by dual-tissue and tumor-only clones. The percentage of dual-tissue clones for each diet-genotype group is displayed at the top of each bar. I) Cell counts of dual-tissue T cell clones identified in the mAT^Tum-adj^. Cell count data is colored in stacked bar graph by course T cell subclusters. J) Cell counts of dual-tissue T cell clones identified in the tumor. Cell count data is colored in stacked bar graph by high-resolution T cell subclusters. K) Cell counts of adipose-only clones identified in the mAT^Tum-adj^. Cell count data is colored in stacked bar graph by course T cell subclusters. L) Cell counts of tumor-only clones identified in the mAT^Tum-adj^. Cell count data is colored in stacked bar graph by high-resolution T cell subclusters. Figure 7A was created with Biorender.com.

Subsequently, within the mAT^Tum-adj^, we quantified the number of adipose-only clonotypes and dual-tissue clonotypes. This analysis revealed that in nearly every group, at least 40% of the clonotypes identified in the mAT^Tum-adj^ were also found in the tumor (Figure 7G). Importantly, this analysis also showed an increase in dual-tissue clonotypes in the lean and obese *Trem2*^-/-^ mAT^Tum-adj^ compared to the diet-matched *Trem2*^+/+^ controls, along with a striking reduction in the number of clonotypes in both weight loss groups. Quantification of clonotypes in the tumor highlighted that a significantly smaller proportion of the clonotypes identified in the tumor were also detected in the mAT^Tum-adj^ (Fig. 7H). Thereafter, we quantified the cell counts for the clonotypes within each tissue and further categorized the data using coarse T cell clustering in the mAT^Tum-adj^ and fine T cell clustering in the tumor. In the mAT^Tum-adj^, the dual-tissue CD8^+^ T cells were increased in the lean *Trem2*^-/-^ samples compared to the lean *Trem2*^+/+^ controls and even further increased in the obese *Trem2*^-/-^ animals. In contrast, the number of CD8^+^ T cells in weight loss *Trem2*^-/-^ mAT^Tum-adj^ decreased compared to the weight loss *Trem2*^+/+^ mAT (Fig. 7I). Concurrently, in the tumor, the cell counts of dual-tissue CD8^+^ TEM, CD8^+^ TEX, and CD4^+^ Th1 cells demonstrated a striking increase in the lean and obese *Trem2*^-/-^ tumors compared to the diet-matched *Trem2*^+/+^ tumors with greatest number of dual-tissue T cells in the lean *Trem2*^-/-^ tumors (Fig. 7J). Similar to the mAT^Tum-adj^, the weight loss tumors exhibited a reduction of dual-tissue T cells in the *Trem2*^-/-^ animals (Fig. 7J). The adipose-only clone cell counts are predominantly composed of NKT and Treg clones, indicating that NKT and Treg populations were mainly restricted to the mAT^Tum-adj^. In contrast, CD8^+^ T cells appear more likely to be identified as a dual-tissue clone between the mAT^Tum-adj^ and the tumor (Fig. 7K). Contrastingly, each diet-genotype group possessed all T cell populations in the tumor-only clone cell counts analysis (Fig. 7L).

### Obese mice are resistant to αPD-1 treatment and do not exhibit an increase in activated T cells

Clinical studies in melanoma and non-small cell lung cancer have suggested obesity may induce a “paradoxical effect” with immune checkpoint inhibitor treatment − a scenario in which obese patients respond more favorably than lean patients^42,43^; however, this has not been well studied in postmenopausal breast cancer or in the context of weight loss. VDJ sequencing and scRNAseq analysis revealed that tumors from *Trem2*^+/+^ lean and weight loss mice had a higher number of all T cell populations per gram tumor and a greater abundance of highly clonal CD8^+^ TEX cells compared to tumors from obese mice (Fig. 6A & J). Consequently, to investigate the response of obese and weight loss mice to αPD-1 treatment in a postmenopausal breast cancer model, ovariectomized mice were placed on LFD and HFD to generate lean, obese, and weight loss groups with food intake and body weight recorded (Supp. Fig. 5A - C). Mice were injected with E0771^PD-L1^ cells at 20 weeks on diet and treated with IgG or αPD-1 antibody during the final 8 d of tumor monitoring (Fig. 8A). Lean and weight loss mice responded to the αPD-1 treatment as evidenced by decreased tumor mass and tumor volume (Fig. 8B, C, E; individual tumor growth curves - Supp. Fig. 5D). Obese IgG and αPD-1 treated groups exhibited no statistical differences in tumor mass or tumor volume at any time point (Fig. 8B & D; Supp. Fig. 5D). To better understand the interactions between the tumor and surrounding mAT, CD11b^+^ cells and T cells were isolated from the stromal vascular fraction of the mAT^Tum-adj^ by fractionation and sequential magnetic sorting using CD11b microbeads followed by CD4/CD8 microbeads (Fig. 8F; Supp. Fig. 5E & F). Gene expression of T cell activation markers *Pdcd1*, *Tigit*, *Tox*, and *Lag3* were increased in the lean αPD-1 mAT^Tum-adj^ T cells compared to the lean IgG (Fig. 8G). Within the obese group, there were no differences in T cell activation marker gene expression between antibody treatment groups; however, *Pdcd1*, *Tox*, and *Lag3* expression were reduced in the obese αPD-1 group compared to the lean αPD-1 treated animals (Fig. 8G). Gene expression on tumor whole tissue lysates revealed an increase in *Cd8a*, *Pdcd1*, *Lag3*, as well as *Trem2* in the weight loss αPD-1 treatment group with no differences found in the obese group (Fig. 8H). In addition, expression of LAM genes *Trem2*, *Plin2*, and *Cd36* in the isolated CD11b^+^ cells reproduced an increase of LAMs in obese mAT^Tum-adj^ and mAT^Contra^ that was not impacted by αPD-1 treatment (Supp. Fig. 5G & H).

**Figure 8.**
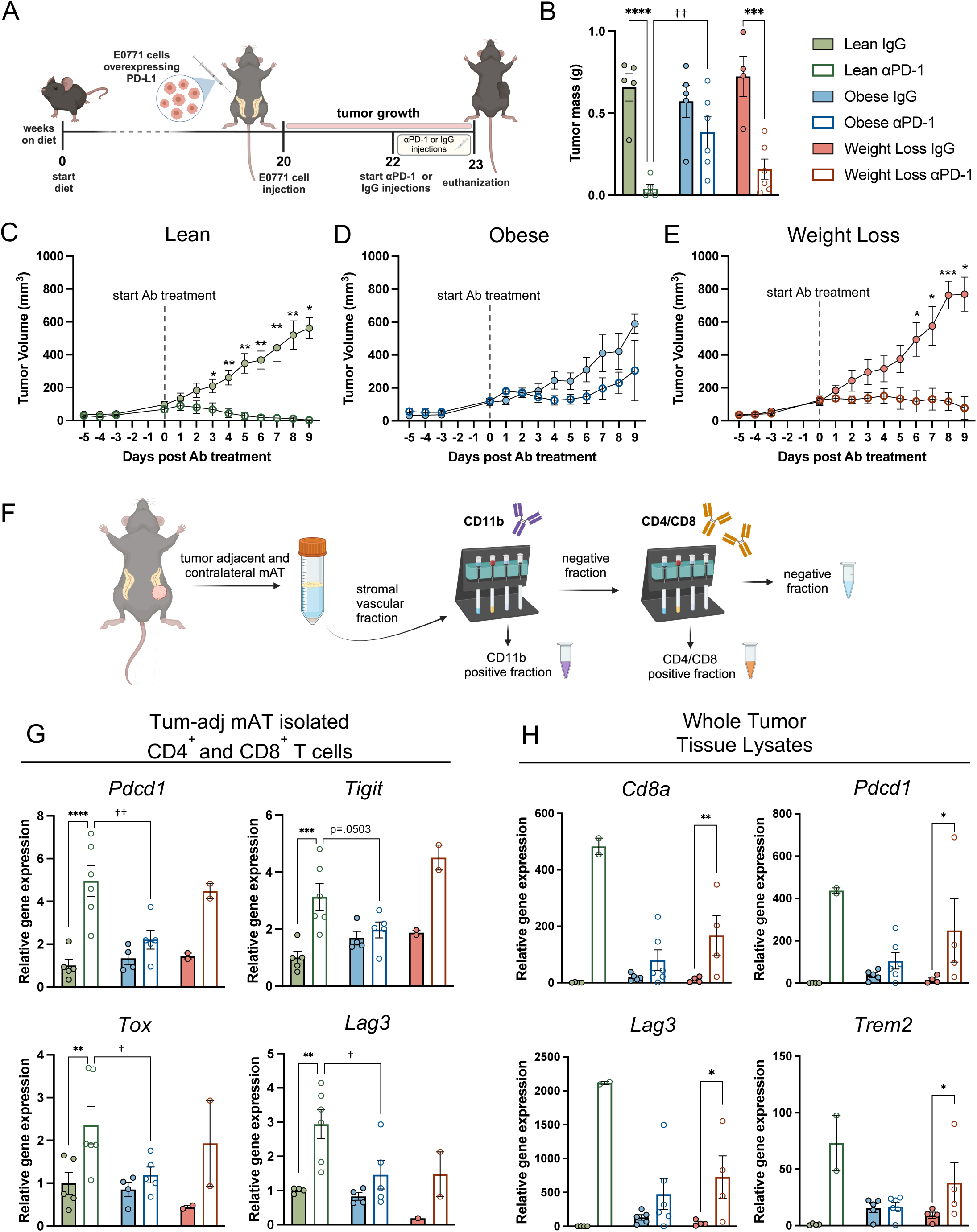
Obese mice do not respond to αPD-1 therapy and do not exhibit an increase of T cell activation markers in tumor-adjacent mammary adipose tissue (mAT^Tum-adj^) or tumor. A) Schematic of study design for lean, obese, and weight loss breast cancer model treated with IgG and αPD-1. During the last two weeks of tumor monitoring, mice were injected with either αPD-1 or IgG isotype control. B) Tumor mass (g) at study endpoint. C-E) Tumor volume (mm^3^) over time for lean, obese, and weight loss IgG and αPD-1 treated mice with dashed line to indicate start of antibody treatment. F) Schematic of isolating the stromal vascular fraction from the mAT^Tum-adj^ and subsequently isolating a pooled sample of CD4^+^ and CD8^+^ T cells using the magnetic Miltenyi Octomacs microbead sorting system. G) T cell activation markers, *Pdcd1*, *Tigit*, *Tox*, and *Lag3* relative gene expression by RT-PCR from magnetically sorted CD4^+^ and CD8^+^ T cells from the mAT^Tum-adj^. H) *Cd8a*, *Pdcd1*, *Lag3*, and *Trem2* gene expression by RT-PCR from whole tumor lysates. Due to the response to αPD-1 treatment in the lean mice, only two tumors from the lean αPD-1 group could be analyzed by RT-PCR. For diet groups, solid colors: green = lean IgG, blue = obese IgG, pink = weight loss IgG; open circles: dark green = lean αPD-1, dark blue = obese αPD-1, maroon = weight loss αPD-1. Two-way ANOVA with uncorrected Fisher’s LSD test was used to compare groups for tumor mass. Mixed effects analysis was used for statistical analysis of tumor volumes over time. Two-way ANOVA with Sidak’s multiple comparison test was used to compare groups in the T cell activation marker gene expression analysis. All data plotted as mean ± SEM for 2-6 mice per group. *denotes *antibody* effect (IgG vs. αPD-1), †denotes *diet* effect, *p<0.05, **p<0.01, ***p<0.001, ****p<0.0001; †p<0.05, ††p<0.01. Figures 2A, F, & H were created with Biorender.com.

## Discussion

In this study, we developed lean, obese, and weight loss models of postmenopausal breast cancer to evaluate the effects of *Trem2* deficiency and immune checkpoint therapy, resulting in three main insights: 1) *Trem2* deficiency attenuated tumor growth in lean, but not obese or weight loss mice. 2) Identical T cell clonotypes were detected in the tumor and mAT^Tum-adj^ of the same animal. 3) αPD-1 treatment restricted tumor growth of PD-L1-expressing tumor cells in lean and weight loss mice, but obese mice demonstrated resistance to immune checkpoint therapy. Overall, our study contributes to a deeper understanding of how weight history impacts the immunological landscape of postmenopausal breast cancer and the potential for TREM2 targeting, with implications for personalized medicine.

Our study, along with previous research in colorectal cancer, sarcoma models, breast cancer, and ovarian cancer^15,17,18^, confirms that *Trem2* deficiency inhibits tumor growth in chow-fed mice. ScRNAseq of tumor immune cells under *Trem2*-deficient conditions by other groups has revealed an increase in CD8^+^ T cells in the MCA/1956 sarcoma model^15,17^ and an increase in both CD8^+^ T cells and NKT cells in the PY8119 TNBC model^44^. In agreement with these findings, we also report an increase in CD4^+^ Th1 and CD8^+^ TEM cells in tumors from lean *Trem2*^-/-^ mice. However, previous studies by other groups were limited to the lean, chow-fed state. Our study focused on human-relevant conditions by examining the effects of *Trem2* deficiency in both obese and weight loss conditions. In addition, we assessed T cells and T cell clonality. In lean *Trem2*^+/+^ tumors, the predominant clonal populations are exhausted CD8^+^ T cells; however, in lean *Trem2*^-/-^ tumors, these populations shift toward an effector memory state and is also associated with an increase of CD4^+^ Th1 cells. This shift does not occur in obese or weight loss states, where the clonal cells predominantly remain exhausted. Greater numbers of TEM cells suggests a potent anti-tumor immune response occurred that was supported by the high frequency of CD4^+^ Th1 cells driving a pro-inflammatory response. Ultimately, this unique cellular composition could contribute to the tumor attenuation phenotype found in lean, but not obese or weight loss, states.

By utilizing scRNAseq and VDJ sequencing in both the tumor and mAT^Tum-adj^, we were able perform side-by-side analysis of T cell clonotypes in these two tissues from the same animal. To our knowledge, this approach has not been performed previously for any cancer types. Intriguingly, we found numerous incidents of identical T cell clonotypes within the tumor and mAT^Tum-adj^ of the same mouse, which we termed “dual-tissue clones”. Notably, in the mAT^Tum-adj^, 40% or more of the T cell clonotypes were dual-tissue clones in nearly all conditions, even reaching 74% in the lean *Trem2*^-/-^ tissue. In fact, the number of dual tissue clones nearly doubles in lean *Trem2*^-/-^ compared with lean *Trem2*^+/+^ mice, suggesting that absence of Trem2 on myeloid cells has immediate consequences for clonal expansion of T cells. This effect of Trem2 deficiency was also seen in obese, but not weight loss mice. The tumor contains an overwhelming number of T cells compared to the mAT^Tum-adj^; however, up to 4.1% of tumor clonotypes were identified as dual-tissue, with thousands of dual-tissue clonal cells detected in the tumor. Interestingly, tumor dual-tissue clone counts were reduced in the obese tumors and nearly ablated in weight loss, suggesting some perturbation resulting in loss of dual-tissue clones could have occurred in obesity that were not recovered with weight loss. Overall, these data suggest the potential for trafficking of T cells between tumors and nearby adipose tissue. This observation could indicate that the adipose tissue may even act as a reservoir of T cells for the tumor. The absence of tumor attenuation in weight loss *Trem2*^-/-^ mice, coupled with the near ablation of dual-tissue clones in the tumor and mAT^Tum-adj^, suggests that the potential trafficking of dual-tissue clones is a crucial component of anti-tumor immunity associated with blocking Trem2. The minimal occurrence of identical clonotypes between the tumor and mAT^Tum-adj^ that were not derived from the same mouse (data not shown) strengthens our confidence in the data, further supporting the trafficking pattern of cells between the two tissues. T cell clonality in tumor-associated adipose tissue may also have relevance for other cancers where obesity serves as a prognostic indicator, including colorectal carcinoma and pancreatic cancer. In the future, this finding could have implications for developing treatments to invigorate adipose tissue T cells against tumors.

Despite the known association between obesity and cancer risk, current research reveals that in certain types of cancer, obesity may improve patient response to immune checkpoint inhibitor treatment. This phenomenon has been termed the “obesity paradox”. Retrospective studies including non-small cell lung cancer^43^, melanoma^42^, and renal cell carcinoma^45^ have demonstrated improved overall survival and progression-free survival for obese patients treated with αPD-1 or αPD-L1 therapy. A meta-analysis including 16 cancer types demonstrated improved overall survival and progression free survival for obese breast cancer patients compared to patients with a BMI < 30^46^. However, for breast cancer, this study only included 8 obese patients and did not indicate menopausal status, which highlights the need for further research on breast cancer and immune checkpoint therapy within the context of obesity. Clinical trial data assessed by Pietenpol’s lab demonstrated obese TNBC patients trended towards increased benefit from combination therapy of carboplatin and atezolizumab, an αPD-L1 therapy, compared to normal and overweight patients^47^. To our knowledge, no retrospective human studies have investigated the interplay between weight loss and immune checkpoint inhibitors in breast cancer, leaving a significant gap in our understanding of this relationship. In our studies, we show that the postmenopausal obese mice do not respond to αPD-1 therapy, which contradicts the obesity paradox. Previous work from the Makowski lab showed that pre-menopausal obese mice responded to the αPD-1 therapy^48^. Other differences between these studies are PD-L1 expression on the tumor cells in our study, and the use of irradiated diets in the Makowski study. Interestingly, Makowski also shows that premenopausal obese mice do not respond to αPD-L1 treatment^22^. In human studies, the level of PD-L1 expression on the tumor cells as well as the menopausal status may be important contributors to whether the obesity paradox prevails. Gaining insights into the interplay of these variables combined with obesity and weight loss will be essential for improving breast cancer patient outcomes.

The effectiveness of PD-1 blockade therapies is thought to hinge upon the ability to reinvigorate progenitor CD8^+^ TEX cells, which have the potential to regain their effector functions, unlike terminally exhausted CD8^+^ T cells^49^. Some studies have suggested that the excess adiposity in obesity induces an increased population of PD-1^+^ CD8^+^ T cells, which has been implicated in the obesity paradox^50^. Analysis of the *Trem2*^+/+^ groups in the scRNAseq data from our study reveals a significant decrease in the number of total CD8^+^ T cells per gram tumor and a reduction in highly clonal CD8^+^ T cells in obese tumors compared to lean and weight loss conditions. The scarcity of T cells in obese tumors may account the absence of a response to αPD-1 therapy. It is also possible that CD8^+^ TEX cells in obese tumors are terminally exhausted, whereas the CD8^+^ TEX in lean and weight loss tumors might not yet be terminally exhausted, thus permitting efficacy of αPD-1 treatment. Further studies are required to assess how breast tumors in lean, obese, and weight loss settings respond to αPD-1 therapy, including retrospective studies on breast cancer patients. Future findings from these studies should guide clinicians to consider weight history when evaluating immune checkpoint therapies, recognizing that responses of different types of tumors may vary in the obese context.

Recently, weight loss drugs, especially GLP-1 receptor agonists, have surged in popularity due to their efficacy and transformative impact on weight management, highlighting the importance of addressing weight loss in our studies. Considering that our study indicates tumors respond differently under weight loss conditions compared to a lean state, it is crucial for clinicians to consider patients’ weight history when considering treatment options – especially when immunotherapies will be employed. Furthermore, many patients are prescribed GLP-1 receptor agonists, and after losing weight, discontinue use of the medication and subsequently regain the weight. Given the rise in weight loss-associated medications, and our limited knowledge on the implications of these therapeutics on tumor growth, future studies must continue to investigate both weight loss and weight regain in the context of both weight loss-associated medications and caloric restriction.

A key strength of our study is that all diet-genotype groups in the scRNAseq and VDJ sequencing experiments included reasonable sample sizes of 3 or 4 per group, with samples processed on separate lanes, ensuring independent and robust biological replication. A limitation of our study is that while the identification of identical T cell clonotypes in the tumor and mAT^Tum-adj^ suggests trafficking of T cells between the two tissues, our data does not explicitly show the movement of the T cells. Accordingly, while we hypothesize that the adipose tissue could be acting as a reservoir of T cells for the tumor, we are not able to show any directionality of any potential trafficking, which means the T cells could also be moving from the tumor to the surrounding adipose tissue. Finally, since scRNAseq and VDJ sequencing only analyze a subset of the immune cells from the tumor and mAT^Tum-adj^, our interpretation of the data is limited to drawing inferences from the cells that were captured. Future studies from this work should include analysis of lean, obese, and weight loss response to immune checkpoint therapy in the context of *Trem2* deficiency or antibody treatment. Additionally, although many patients are either obese or have lost weight, weight cycling – characterized by alternating periods of weight gain and loss^10^ – is more common and should be addressed in future studies.

By examining obesity and weight loss across various contexts, including *Trem2* deficiency and immune checkpoint therapy, and assessing both the tumor and surrounding adipose tissue in all studies, we have gained significant insights into how weight history affects both adipose tissue and the tumor microenvironment. Our data suggest a potential crosstalk between these tissues, possibly involving T cell trafficking. These findings indicate that weight history is a critical factor for clinicians to consider in patient care, as its effects may vary depending on the specific clinical scenario.

## Funding

This project was funded by a Pilot and Feasibility Award from the Vanderbilt Ingram Cancer Center (VICC; P30CA068485) Breast Specialized Program of Research Excellence (SPORE; P50CA098131) program to AHH and a Mark Foundation Endeavor Award to JCR, LM, and KEW. EWP was supported by the Immunological Mechanisms of Disease Training Program (5T32Al138932) and the Mark Foundation Endeavor Award. AHH was supported by a Career Scientist Award from the Veterans Affairs (IK6 BX005649) and a Veterans Affairs Merit Award (I01 BX002195). LM and JCR are supported by NCI U01CA272541 and LM by R01CA253329. BDL is supported by a Department of Defense Breast Cancer Research Program grant (BC201286). JAP is supported by a Breast SPORE (P50CA098131).

## Author’s Information

AHH is now Vice-Provost and Senior Associate Dean for Faculty Affairs and Career Development at University of Texas Southwestern in Dallas, TX

## Acknowledgments

The authors would like to thank Jamie Garcia, PhD, for assistance with cell isolation and preparation for scRNAseq. Ovariectomies were performed at the VMMPC, and the authors would like to thank Carlo Malabanan, Alicia Kellarakos, Tiffany Farmer, and Teri Doss for performing the ovariectomy surgeries. Tumor cell injections and body composition measurements were also performed in the VMMPC, which is supported by DK135073 (MMPC-Live) and DK020593 (DRTC). Isolation of CD45^+^ cells in preparation for scRNAseq was performed in the VUMC Flow Cytometry Shared Resource. The VUMC Flow Cytometry Shared Resource is supported by the Vanderbilt Ingram Cancer Center (P30 CA68485) and the Vanderbilt Digestive Disease Research Center (DK058404). scRNAseq and VDJ sequencing was performed in the Vanderbilt Technologies for Advanced Genomics (VANTAGE) core laboratory, which is supported in part by Clinical and Translational Science Award Grant 5UL1 RR024975-03, Vanderbilt Ingram Cancer Center Grant P30 CA68485, Vanderbilt Vision Center Grant P30 EY08126, and National Institutes of Health/National Center for Research Resources Grant G20 RR030956. Analysis of scRNAseq and VDJ sequencing utilized the resources provided by the Vanderbilt Advanced Computing Center for Research and Education (ACCRE), which is operated by and for Vanderbilt faculty.

