## Supplemental Figures for "Trem2 deficiency attenuates breast cancer tumor growth in lean, but not obese or weight loss, mice and is associated with alterations of clonal T cell populations"

### Supplemental Figure 1

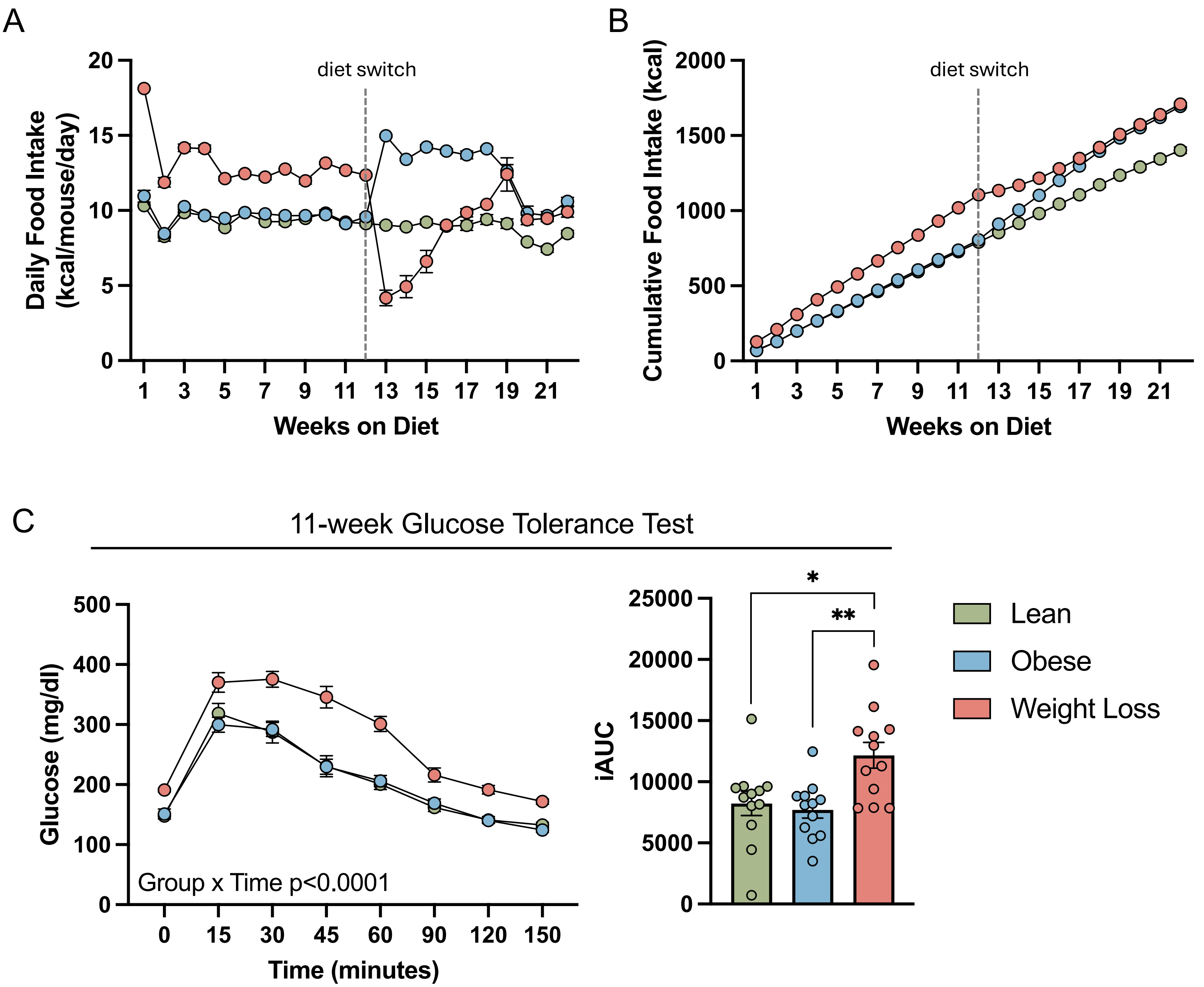


**Supplemental Figure 1. Food intake and pre-diet switch glucose tolerance in lean, obese, and weight loss mice.**

Following ovariectomy, mice were placed on 10% low fat diet (LFD) and 60% high fat diet (HFD) as shown in Figure 1A. A) Mouse daily food intake represented as kcal/mouse/day calculated as an average over the week with dashed line indicating diet switch. B) Mouse cumulative food intake (kcal), with dashed line indicating diet switch. C) Blood glucose (mg/dL) measured over time during 11-week time point ipGTT and corresponding iAUC. For diet groups, green = lean, blue = obese, pink = weight loss. Repeated measures two-way ANOVA was used for statistical analysis of blood glucose over time in ipGTT. One-way ANOVA with Tukey's multiple comparisons test was used to compare groups for iAUC. All data plotted as mean ± SEM for 12 mice per group. *p<0.05, **p<0.01.

### Supplemental Figure 2

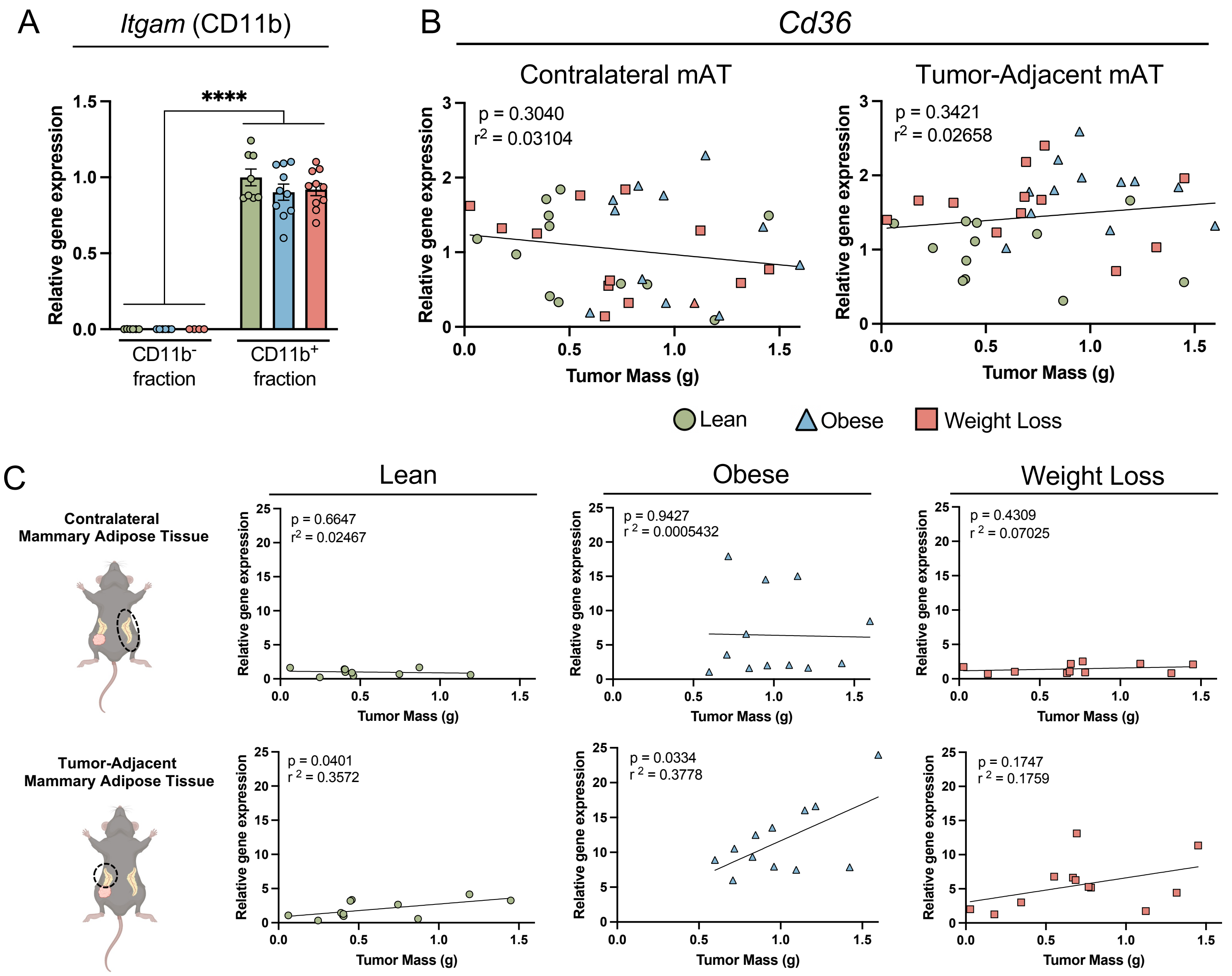


**Supplemental Figure 2. Mammary adipose tissue lipid associated macrophage marker gene expression and correlation with tumor mass.**

A) *Itgam* (CD11b) relative gene expression by RT-PCR in $CD11b^-$ and $CD11b^+$ tumor fractions. B) Tumor mass vs. *Cd36* relative gene expression in contralateral and tumor-adjacent mammary adipose tissue (mAT) with linear regression fit. C) Tumor mass vs. *Trem2* relative gene expression in contralateral and tumor-adjacent mAT separated by diet group and adipose tissue depot with linear regression fit. Two-way ANOVA with Tukey's multiple comparison tests were used to compare groups for *Itgam* gene expression. Simple linear regression analysis was applied to correlation plot. All data plotted as mean ± SEM for 10-12 mice per group. ****$p<0.0001$. Statistical values for correlation analysis indicated on graphs. Supplemental Figures C was created with Biorender.com.

### Supplemental Figure 3

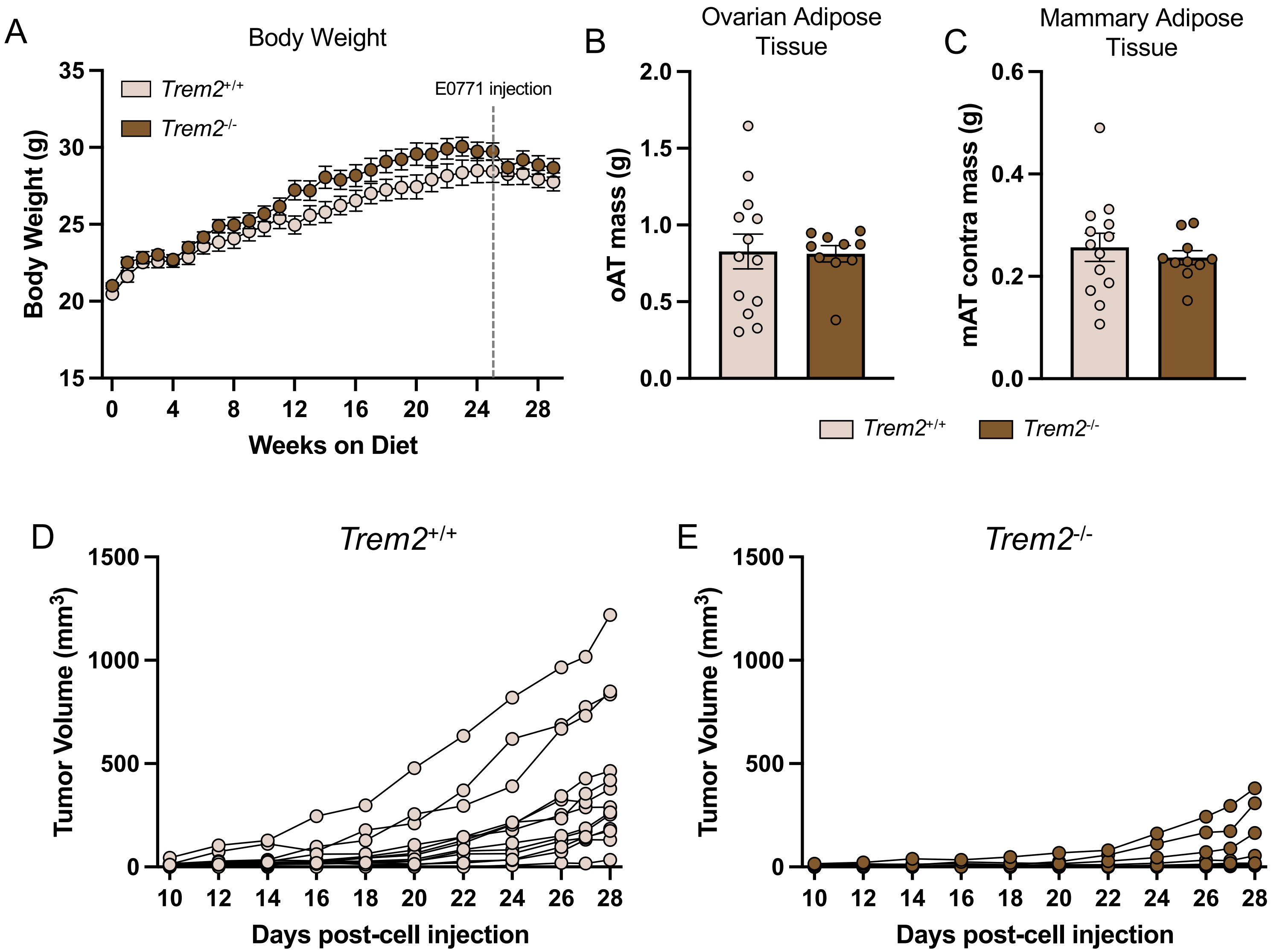


**Supplemental Figure 3. Individual tumor volumes over time and oAT and mAT mass at study endpoint.**
A) Body weights (g) measured weekly with dashed lines to indicate E0771$^{PD-L1}$ cell injection. B) Ovarian adipose tissue mass (g) at study endpoint. C) Mammary adipose tissue mass (g) at study endpoint. D) Tumor volumes (mm$^3$) over time for Trem2$^{+/+}$ mice. E) Tumor volumes (mm$^3$) over time for Trem2$^{-/-}$ mice. Unpaired two-tailed t tests were used to compare genotypes for ovarian and mammary adipose tissue mass. All data plotted as mean ± SEM for n=10-13 per group. No data indicated as statistically significant.

### Supplemental Figure 4

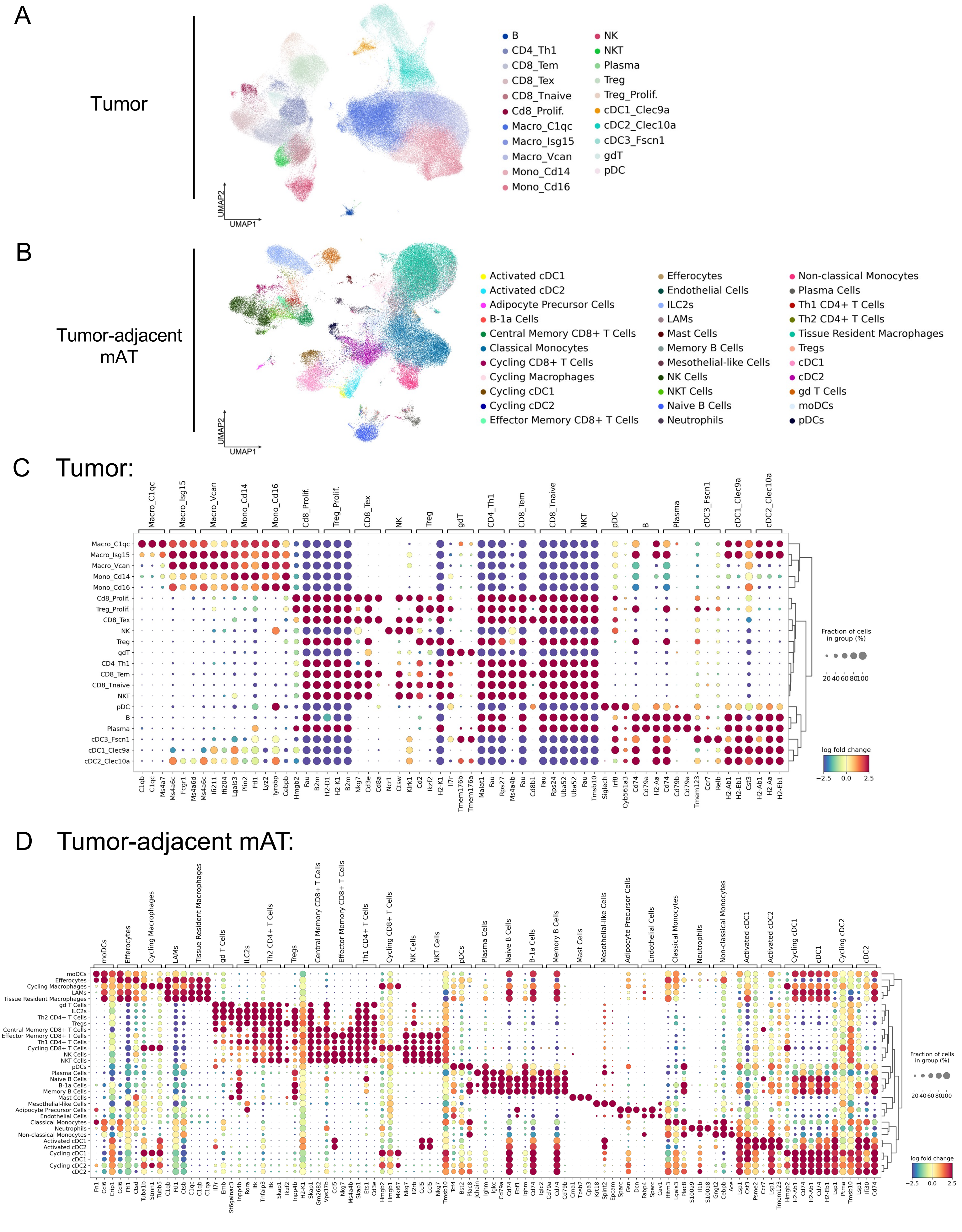

**Supplemental Figure 4. Immune cell populations in tumors and tumor-adjacent mammary adipose tissue (mAT) analyzed by single-cell RNA sequencing.**

A-B) Unbiased clustering of 201,912 single cells in the tumor (A) and 98,532 single cells in the tumor-adjacent mAT (B) colored by high-resolution cell type identities via Uniform Manifold Approximation and Projection (UMAP). C-D) Dot plot of the top 3 differentially expressed genes (DEGs) for all high resolution clusters in the tumor (C) and tumor-adjacent mAT (D) with dot size representing the proportion of cells expressing the gene and color indicating the log fold change.

### Supplemental Figure 5

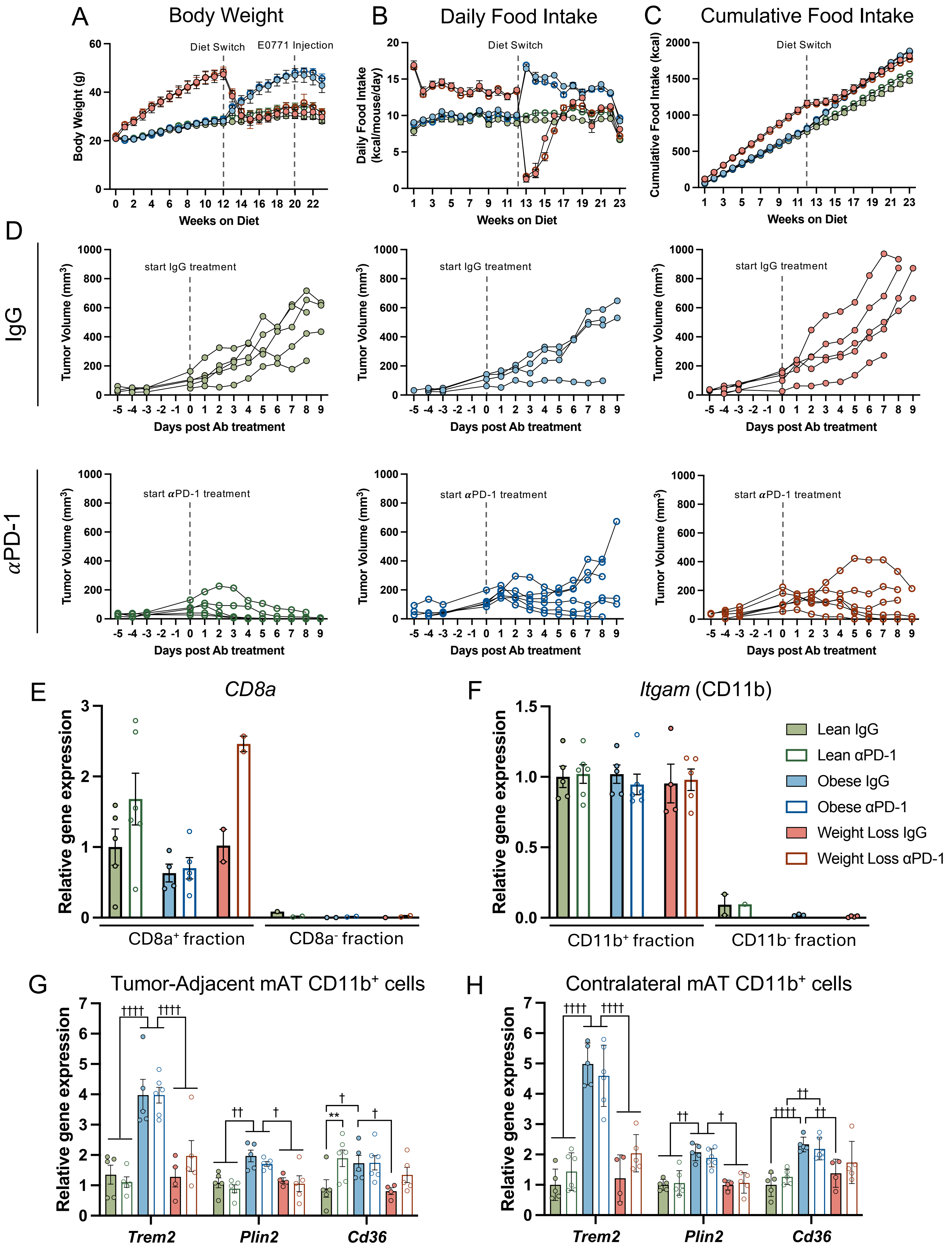

**Supplemental Figure 5. IgG and αPD-1 treatment in lean, obese, and weight loss mice.**
A) Body weights (g) for all groups measured weekly with dashed lines to indicate diet switch and E0771 cell injection. B) Mouse daily food intake represented as kcal/mouse/day calculated as an average over the week with dashed line indicating diet switch. C) Mouse cumulative food intake (kcal) over the entire time on diet with dashed line indicating diet switch. D) Tumor volume ($mm^3$) over time for all individual tumors with dashed line to indicate start of IgG or αPD-1 treatment. Data separated by diet and treatment group. E) *Cd8a* relative gene expression by RT-PCR in $CD4^+/CD8^+$ and $CD4^-/CD8^-$ tumor-adjacent mAT fractions. F) *Itgam* (CD11b) relative gene expression by RT-PCR in $CD11b^+$ and $CD11b^-$ contralateral mAT fractions. G) Relative gene expression by RT-PCR of LAM genes *Trem2*, *Plin2*, and *Cd36* from the tumor-adjacent mAT $CD11b^+$ fraction. H) Relative gene expression by RT-PCR of LAM genes *Trem2*, *Plin2*, and *Cd36* from the contralateral mAT $CD11b^+$ fraction. For diet groups, solid colors: green = lean IgG, blue = obese IgG, pink = weight loss IgG; open circles: dark green = lean αPD-1, dark blue = obese αPD-1, maroon = weight loss αPD-1. Two-way ANOVA with uncorrected Fisher's LSD test was performed to compare groups for all LAM gene expression. All data plotted as mean ± SEM of 2-6 mice per group. *denotes *antibody* effect (IgG vs. αPD-1), †denotes *diet* effect, **$p<0.01$; †$p<0.05$, ††$p<0.01$, ††††$p<0.0001$.
